# Data science competition for cross-site delineation and classification of individual trees from airborne remote sensing data

**DOI:** 10.1101/2021.08.06.453503

**Authors:** Sarah Jane Graves, Sergio Marconi, Dylan Stewart, Ira Harmon, Ben G. Weinstein, Yuzi Kanazawa, Victoria M Scholl, Maxwell B Joseph, Joseph McClinchy, Luke Browne, Megan K Sullivan, Sergio Estrada-Villegas, Eduardo Tusa, Daisy Zhe Wang, Aditya Singh, Stephanie A Bohlman, Alina Zare, Ethan P. White

## Abstract

Delineating and classifying individual trees in remote sensing data is challenging. Many tree crown delineation methods have difficulty in closed-canopy forests and do not leverage multiple datasets. Methods to classify individual species are often accurate for common species, but perform poorly for less common species and when applied to new sites. We ran a data science competition to help identify effective methods for delineation of individual crowns and classification to determine species identity. This competition included data from multiple sites to assess the methods’ ability to generalize learning across multiple sites simultaneously, and transfer learning to novel sites where the methods were not trained. Six teams, representing 4 countries and 9 individual participants, submitted predictions. Methods from a previous competition were also applied and used as the baseline to understand whether the methods are changing and improving over time. The best delineation method was based on an instance segmentation pipeline, closely followed by a Faster R-CNN pipeline, both of which outperformed the baseline method. However, the baseline (based on a growing region algorithm) still performed well as did the Faster R-CNN. All delineation methods generalized well and transferred to novel forests effectively. The best species classification method was based on a two-stage fully connected neural network, which significantly outperformed the baseline (a random forest and Gradient boosting ensemble). The classification methods generalized well, with all teams training their models using multiple sites simultaneously, but the predictions from these trained models generally failed to transfer effectively to a novel site. Classification performance was strongly influenced by the number of field-based species IDs available for training the models, with most methods predicting common species well at the training sites. Classification errors (i.e., species misidentification) were most common between similar species in the same genus and different species that occur in the same habitat. The best methods handled class imbalance well and learned unique spectral features even with limited data. Most methods performed better than baseline in detecting new (untrained) species, especially in the site with no training data. Our experience further shows that data science competitions are useful for comparing different methods through the use of a standardized dataset and set of evaluation criteria, which highlights promising approaches and common challenges, and therefore advances the ecological and remote sensing field as a whole.

## Introduction

High resolution remote sensing imagery provides critical information about the presence and types of organisms within and among ecosystems at scales beyond those observable using field techniques. Generating ecological data on individual organisms from imagery requires the image processing tasks of delineation (Ke & Quackenbush, 2011): the ability to detect, isolate, and outline organisms from background information; and classification (Fassnacht et al., 2016): the ability to assign a meaningful label to the organism. While advancements of these tasks have been made in forest ecology, challenges remain to delineate individual tree crowns and classify them to their taxonomic species, especially in taxonomically and structurally diverse forests with closed canopies (Ke & Quackenbush, 2011; Fassnacht et al., 2016).

Data science competitions are a unique way to advance image processing methods for particular applications (Carpenter, 2011). These competitions provide a standardized dataset and criteria for evaluation, and have the potential to draw expertise from different domains because of the focus on common data science tasks (Dorr et al., 2016; Marconi et al., 2019; Van Etten, Lindenbaum & Bacastow, 2019). While competitions have allowed for the advancement of many applications, ecology is just beginning to use competitions (e.g., Humphries et al., 2018; Little et al., 2020), due in part to the increasing availability of large, openly available ecological datasets (Marconi et al., 2019).

In 2017, we ran a competition that used data from the National Ecological Observatory Network (NEON) to generate species predictions of individual tree crowns in a single temperate forest (Marconi et al., 2019). NEON is a 30-year effort of the National Science Foundation to collect standardized organismal, biogeochemical, and remote sensing data over 81 sites in the US (Schimel et al., 2007). The data provided by NEON covers a broad array of ecosystem components including data on trees and associated airborne remote sensing imagery. NEON data is ideal for use in data science competitions because it is openly available, well documented, and part of a massive continental scale data collection effort. Therefore, methods and lessons learned from the competition can be applied to a large-scale open dataset being used by large numbers of researchers. The 2017 event was the first data science competition using NEON data, and was instrumental in advancing methods and in establishing a framework for providing data and evaluating submissions. The competition identified the most effective methods for delineating individual tree crowns from airborne remote sensing data, aligning delineations to field data, and assigning a species label to delineations. Evaluating the performance and comparing approaches allowed us to identify critical components for success and approaches that produced the best information. The results of the 2017 competition showed that hyperspectral reflectance, which captures biophysical and chemical properties of vegetation, is useful for delineations. Moreover, the results showed that there is a need for improvement in the delineation of small tree crowns. We also learned that most successful approaches to species classification are multi-part, requiring data cleaning to remove noise and outliers prior to analysis, and that methods incorporating uncertainty in predictions show the most promise for future application.

While the first competition was an important step towards better methods for converting remote sensing to ecological information, we need to evaluate how methods work when trained on standard forest inventory data across large spatial scales and diverse forest types, and how well such methods apply when making predictions in novel forests. For delineation, this situation presents unique image processing challenges due to a wide range of crown shapes, canopy complexities, and backgrounds types, caused by the diversity of forest composition and structure. For classification, this task is challenging due to highly imbalanced multi-species datasets, combined with differences in the species present at different sites. Finally, differences in the remote sensing data due to variability in the brightness and shadows caused by differences in sun position, and phenological differences in the vegetation due to the time of year, impact the ability for methods to generalize and transfer patterns. While the tasks of delineation and classification remain critical to the needs of ecologists, an expansion in the number sites and diversity of data is required to effectively achieve these tasks.

To address these needs we ran a new iteration of the IDTReeS competition focused on the tasks of delineation and classification using a new set of data that allowed for within and cross site evaluation. This iteration of the competition uses data from three NEON sites in the Southeastern United States to compare 1) how well methods generalize by incorporating information from different sites, and 2) how well methods transfer to make predictions on sites for which the algorithms have not been trained. Here we present the results of the competition that includes delineation and classification scores from participating teams, details about the methods used, and a discussion of how this competition advances our ability to delineate and classify individual trees using existing inventory and remote sensing data. More details of the approaches taken by one of the teams are included in a companion paper (Scholl et al., 2021).

## Materials & Methods

### Study sites

We used multiple NEON data products from three sites in combination with data collected by members from our team. The three NEON sites in the southeastern United States (Fig. 1) used in this study are part of three separate NEON ecoclimatic domains and represent distinct environmental, geographic, and vegetative characteristics (Thorpe et al., 2016). The Ordway-Swisher Biological Station in Florida (OSBS, Southeastern domain, 03) is a mixed forest of hardwood and conifers, mostly dominated by pines. Talladega National Forest in Alabama (TALL, Ozarks complex domain, 08) is a primarily forest of mixed hardwood and conifers, and also dominated by pines. The pine dominated forests at OSBS and TALL tend to have an open canopy, while hardwood dominated forests at these sites usually have a closed canopy. Mountain Lake Biological Station in Virginia (MLBS, Appalachians and Cumberland Plateau domain, 07) has mainly closed-canopy hardwood forest. In MLBS, pine species are more rare than at OBSB and TALL, and there is a greater abundance and diversity of canopy hardwood species.

**Figure 1.**
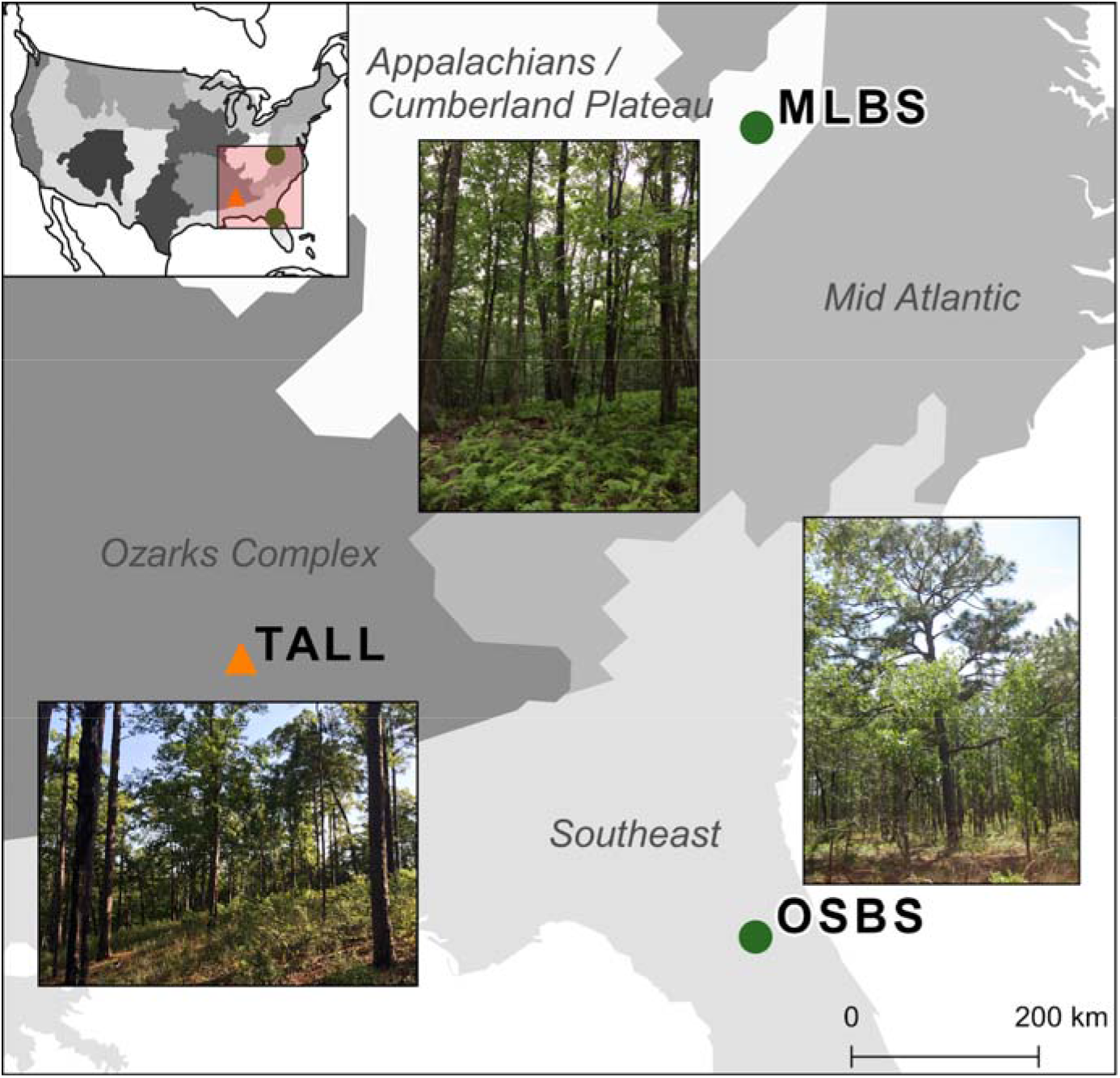
Map of the three study sites and domains of the National Ecological Observatory Network (NEON). The sites are part of three separate NEON ecoclimatic domains and represent distinct environmental, geographic, and vegetative characteristics. Overview map of the USA shows the NEON domains. To evaluate the ability for methods to apply within sites, we used data from the Ordway Swisher Biological Station (OSBS) and Mountain Lake Biological Station (MLBS) for training and testing (green circle). To evaluate the ability for methods to apply to new sites, we used data from the Talladega National Forest (TALL) for testing only (orange triangle).

### NEON data

We use NEON data from two standard collections. The remote sensing data were generated by the NEON Airborne Observation Platform (AOP) and are provided as four different products, each one measuring different properties of the vegetation and the ground surface (Appendix A, Table A1). The AOP data products are high-resolution orthorectified camera imagery (RGB), discrete return LiDAR point clouds (LAS), LiDAR canopy height models (CHM), and spectrometer orthorectified surface directional reflectance - mosaic hyperspectral surface reflectance (HSI). The data products were downloaded using the NEON API. The most recent available years of data are used: 2019 for TALL and OSBS and 2018 for MLBS. Since the data were downloaded from NEON prior to 2021, they are considered Provisional Data.

The field data to provide species labels in the competition training data were collected through the NEON Terrestrial Observation System (TOS). The data contain information on individual tree identifiers, location of trees relative to sampling locations (i.e. distance and azimuth from a central location), species and genus labels, and measures of salient structural attributes. The field attribute that is directly used in this competition is the taxonomic species information. The taxonomic species information is described by its scientific name, which includes a genus and species classification. To simplify the taxonomic species information, each scientific name is simplified to its unique taxonomic identification code. More information about the data products and the field data and the list of species classes and taxonomic codes is provided in Appendix A.

### Individual tree crown data

Individual tree crown (ITC) data are spatial delineations that represent the spatial extent of an individual tree in remote sensing data and are fundamental for both tasks in the competition. For the delineation task, participants were given ITCs in the training dataset and generated ITCs for the test dataset (Fig. 2). For the classification task, participants were given ITCs labeled with the taxonomic species and generated species predictions for unclassified ITCs. ITC delineation data are not a standard NEON product and were generated by the IDTReeS research team. Each ITC is represented by a 2-dimensional rectangular bounding box that geographically defines an individual crown. The delineation represents the maximum crown boundary or extent in the North/South and East/West directions. Because ITC data are time-intensive and difficult to generate, the research team used a combination of two different approaches to produce both a reasonably large number of labeled data for training and precise data for evaluation. Both ITC datasets were generated by experts who are familiar with the ecology of the sites. To generate the training ITCs, the research team in the lab used multiple remote sensing datasets and field inventory data from NEON. To generate the test ITCs, the research team in the field measured tree crown boundaries and identified species. The ITC data were labelled within discrete 20 × 20 m plots. The plots are designated as training and testing for the two tasks as described below. More information about how ITC data are generated and how they are related to the field data from NEON is provided in Appendix A.

**Figure 2.**
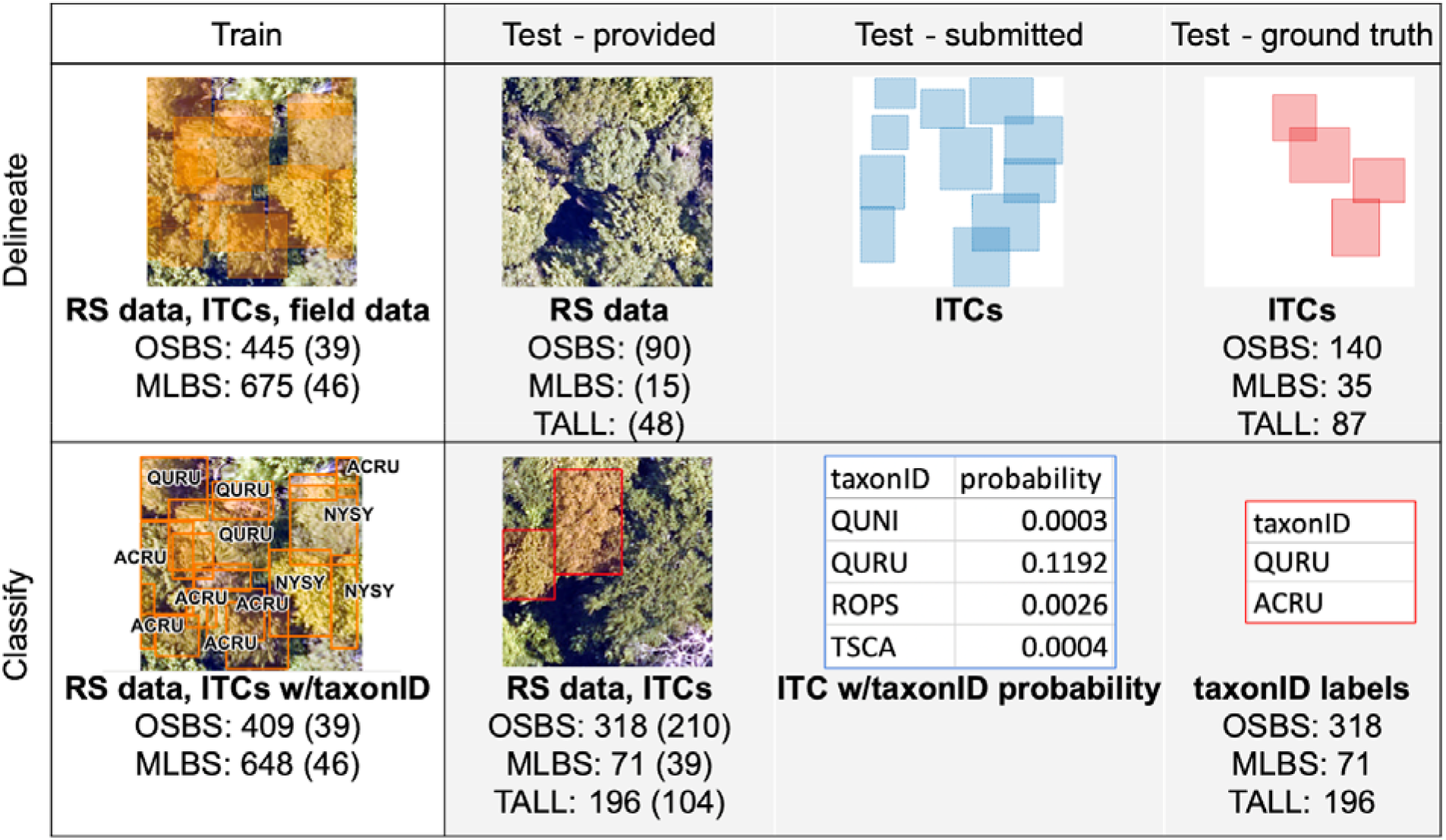
Data used for training and testing of methods for the delineation and classification tasks. Data listed for each of the three sites from the National Ecological Observatory Network (NEON); OSBS = Ordway Swisher Biological Station; MLBS = Mountain Lake Biological Station; TALL = Talladega National Forest. RS data include all four remote sensing data products (Appendix A). ITCs are the number of individual tree crown delineations. The number of plots in the parentheses is the number of 20 × 20 meter plots with associated remote sensing data.

### Solicitation and team participation

The competition was announced on February 3rd, 2020 and advertised to individuals and communities focused on remote sensing, image processing, and forest ecology. We also contacted the 109 people who had registered from the previous competition. In total, there were 130 registrations for this second competition. Submissions were received from 6 participating teams (Appendix B, Table B1). Three teams completed the delineation task and 4 teams completed the classification task. Of the six teams, two completed both tasks.

All teams were allowed up to 4 submissions per task. Submissions made prior to the final submission were evaluated and scores were returned. Pre-submissions were allowed to ensure submissions were properly formatted and provide teams with feedback on model performance. The final submission deadline was extended by 2 months after the train and test data were released. This was done to allow teams more time to work with the data given the challenges associated with COVID-19. The number of pre-submissions was limited to reduce the chance of artificially boosting performance indirectly from the test set. The number of pre-submissions varied by team, with 5 pre-submissions from the Fujitsu and Intellisense_CAU teams (an additional submission allowed due to timeline extension), 4 submissions for Jeepers Treepers, 2 for Más JALApeñoS, and no pre-submissions for INRAE-GIPSA.

### Delineation task

The delineation task required the detection of ITCs, and defining their boundaries using remote sensing data. Participants were given remote sensing data and submitted spatial vector data that defined the location and size of all predicted ITCs. The data were split into training and testing datasets (Fig. 2). Training data were provided for the development and self-evaluation of methods and made available to participants at the start of the competition. Training data included all remote sensing data products (clips of 20 × 20 meters around each plot) for 85 plots of the OSBS and MLBS sites and 1120 ITC delineations for all trees within each plot at OSBS and MLBS. Field measurements were also available for each ITC delineation that provides information on the species, size, and other salient attributes of individuals (see Appendix A). Participants could use any remote sensing and field data for training their methods. No TALL data were provided in the training data. The testing data were separate plots with associated remote sensing data and ITC delineations. Only the remote sensing data for the 153 test plots was provided to the participants. The ITC delineations for the test plots, which were labelled in the field, were withheld from the teams. Participants submitted delineated ITCs for all crowns for each plot. A total of 262 ground truth ITCs across all three sites (a subset of 1-9 trees per plot) were used by the competition organizers to score the team submissions.

We evaluated each possible pair of predicted ITC delineation and ground-truth delineations using two metrics: the Intersection over Union (IoU) and a modification of the Rand index we refer to as RandCrowns (Stewart et al., 2021). The IoU metric, known also as the Jaccard Index (Jaccard, 1908), is defined as the area of overlap between the ground-truth ITC delineation and predicted ITC delineation divided by the total area of both ITC delineations. The RandCrowns measure is designed to provide robust detection scores despite spatial uncertainty in ground truth and label data. RandCrowns is a modification of the commonly used Rand index using a set of “halos” around each ground truth box to account for variability in the size of the ITC delineations with respect to ground-truth. The halos account for variability in the crown size and uncertainty of ground-truth ITC delineations. Scores for both IoU and RandCrown range from 0 (no overlap) to 1 (perfect overlap), but in general, scores are higher for RandCrowns than IoU.

Since the test data included only a subset of ITC delineations for all trees in the plot and participants submitted all possible ITC delineations, the first step was assigning a single predicted ITC delineation to each ground-truth ITC delineation in the test data. This was an important step since participant submissions may include many delineations that overlap the test ITCs. In cases of ambiguity caused by multiple predictions corresponding to a single test ITC delineation, we used the one-to-one mapping that provides the best RandCrowns score for the submission, which was done using the Hungarian assignment algorithm. Each possible pair of predicted and ground-truth delineations was scored using the RandCrowns measure and the highest scoring prediction for each ground-truth delineation was used for evaluation. The IoU measure was computed on each pair of submitted and ground-truth ITCs. For each metric, the mean of all assigned pairs was calculated to produce the final evaluation scores for participants.

### Delineation algorithms

Four unique approaches were used for the delineation task (Appendix B). The baseline delineation method was the approach that performed the best in the first iteration on a single site (Dalponte, Frizzera & Gianelle, 2019) and is available in the itcSegment R package (Dalponte, 2018). It is a region growing method that identifies seed points in a single HSI band (at ∼ 810 nm) using a moving window to identify high reflectance associated with the tallest parts of a tree crown and grows the ITC based reflectance differences of neighboring pixels. Parameters used in the first iteration were implemented rather than optimizing parameters using the training data. The Fujitsu Satellite team used an instance segmentation pipeline. First, the spatial resolution of the HSI data was increased using the RGB images and bands were iteratively selected based on the best performance. Second, a two-branch backbone structure neural network was used to extract image features. These features were then fused and integrated into network, regression, and segmentation modules. The CHM was used to filter delineations based on a 2 m height threshold. The Intellisense CAU team trained a Faster-RCNN detector with a ResNet 50 backbone on the RGB data. A test-time augmentation was implemented to improve the delineations. The INRAE-GISPA team implemented an adaptive 3D mean shift method using the LiDAR point cloud data (Tusa et al., 2020). Critical to this approach was the use of a probability density function (PDF) to identify clusters of points that define an ITC. The PDF was based on a superellipsoid kernel where the parameters that defined the crown shape and kernel size were based on allometric equations using tree height, crown radius, and crown depth.

### Classification task

The goal of the classification task was to classify ITCs to their taxonomic species. Remote sensing-based species identification over large spatial scales would have major benefits for ecology, but this task is challenging due to low inter-species variance in spectral properties and the highly imbalanced nature of these types of datasets (Graves et al., 2016).

Like the delineation task, the data were split into training and testing datasets where the training data allows for the development and self-evaluation of models and the testing data was used to evaluate the team methods. The characteristics of training data for the classification task were similar to the delineation task. Training data included all remote sensing data products (clips of 20 × 20 meters around each plot) for 85 OSBS and MLBS plots, and ITC delineations with taxonomic species labels for 1057 ITCs (Fig. 2). Participants could use any of the remote sensing and field data for training their models since this represents a common scenario where models are developed using data from inventory plots. No TALL data was provided in the training data. The testing data were 353 separate plots with associated remote sensing data, 585 ITC delineations, and ground truth species labels (withheld from the teams) at the OSBS, MLBS, and TALL sites. Providing the ITC delineations kept this task focused on classification methods rather than having participants also incorporate delineation approaches prior or post classification. Participants submitted the taxonomic species predictions for ITCs in the test data. The predictions were submitted as a probability from 0 to 100% that the ITCs belonged to the associated species class. The ground truth species labels for each ITC were used by the competition organizers to score the submissions.

Significant features of this dataset, and forest remote sensing data in general, are class imbalance in the training data, and a difference in species composition and relative abundances between the training data and the test data. Due to the nature of these data, the ability to train on imbalanced data and predict species with species identities and abundances that differ between the training and testing datasets is an important challenge addressed in this competition. The training dataset for the OSBS and MLBS sites had a total of 33 distinct species classes, ranging from 1-302 individuals per species class (Fig. 3). This distribution represents the composition and relative abundance of canopy trees in the NEON plots and therefore the data available from forest inventory plots that are used to develop and test classification models. The test data for OSBS and MLBS both show unequal distributions of data among species classes. The test data for both sites include only 15 species in the training data, and both sites include species in the test data that are not part of the training data (OSBS: 11 species, MLBS: 5 species). Furthermore, while the test data for TALL has less imbalance across the species classes than the training data at OSBS and MLBS, it includes only 10 of the species from the training data and introduces 11 new species that are not part of the training data (as “Other” in Fig. 3). In this way, the external TALL site tests not only the ability of the models to be applied to new remote sensing data, but also to a new site with different species composition.

**Figure 3.**
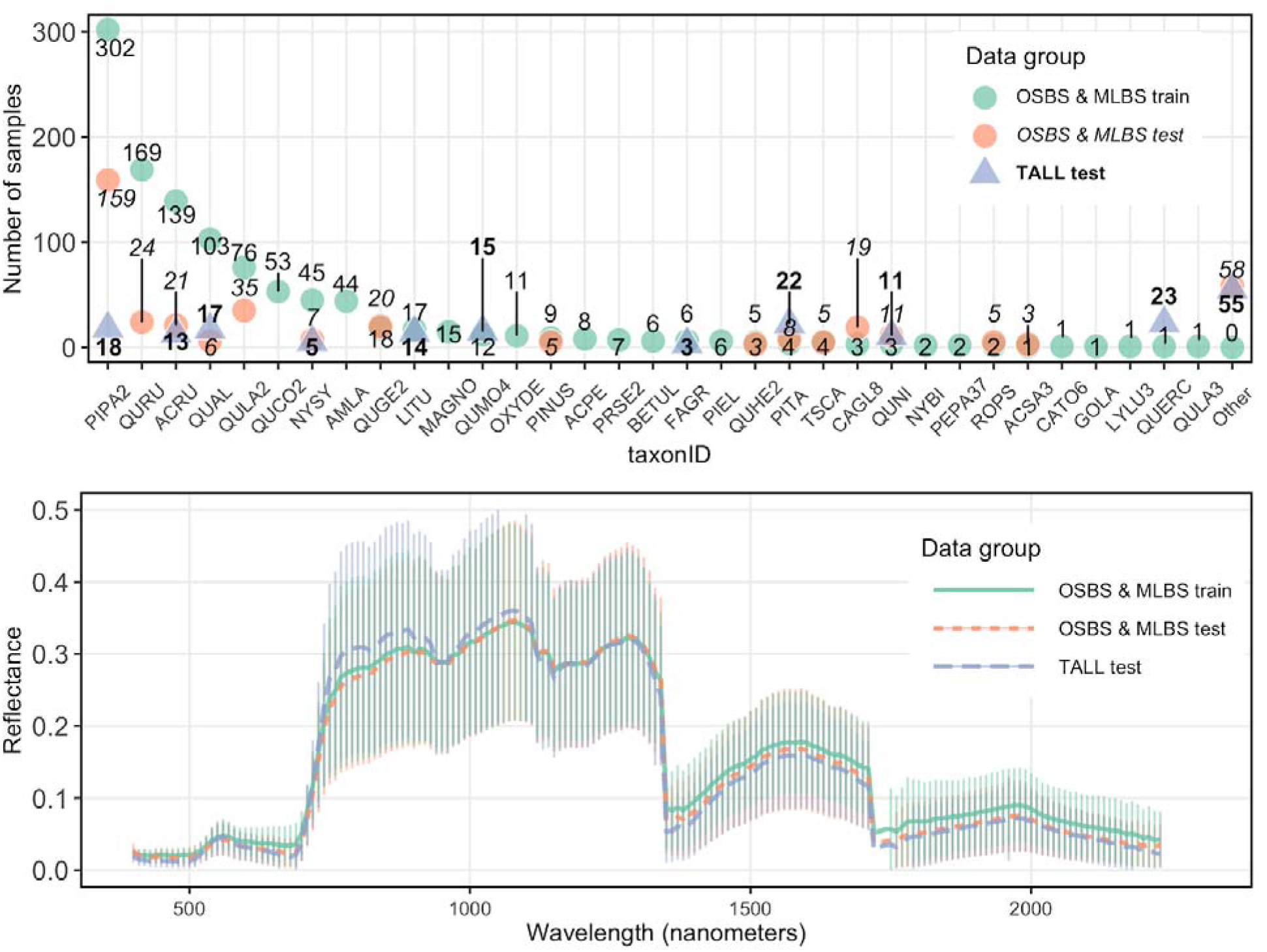
Distribution of samples and reflectance for the classification task. Top: Distribution of samples per species class (taxonID) for each dataset. The number of samples for each taxonomic class differs in training and test data for OSBS and MLBS sites and test data for the TALL site. Taxonomic class is arranged based on the number of data points in the train data. Bottom: Hyperspectral reflectance for each data group. Reflectance sampled from 100 random pixels of 10 random 20 × 20 meter plots for each site in the training and test data. Mean (thick lines) and standard deviation calculated for each data group.

To account for new species classes in the test dataset not present in the training data, participants were also allowed to include a species class with the label “Other”. The Other class can be used to indicate a probability that an ITC is a species that is not represented in the training data and is therefore likely a new species.

### Evaluation of the classification task

The primary metric for evaluation of submissions was the macro F1 score. This score is the arithmetic mean of the weighted harmonic mean of the precision and recall scores for each species class in the dataset and is given by equation (1), where C is the set of species classes, P_c_ is the precision of species class c, R_c_ is the recall of species class c, and |C| is the cardinality of set C.

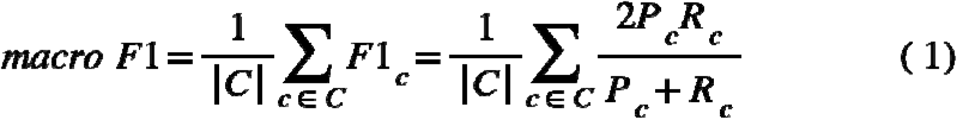

The score weights each species class equally regardless of the number of individuals in each class. When used in combination with other metrics, macro F1 score is a good indicator of classifier bias that may result from training on an imbalanced dataset. Since macro F1 score requires hard classification labels for each ITC and species predictions were submitted as the probability of a species label, the label with the highest probability was selected for each ITC.

We evaluated participant models on two additional metrics, log-loss and weighted F1 score, to better understand their performance. While these metrics were not used to assess the strongest submission, they provide another measure of model robustness and allow for a more in-depth comparison between models. Cross-entropy loss, also known as log loss, is a performance metric calculated from the species class membership probability for each ITC, y-hat, and the probability for each label in the test set, y. Equation (2) gives the formula for log-loss, where N is the number of instances in the test set and k is the number of classes.

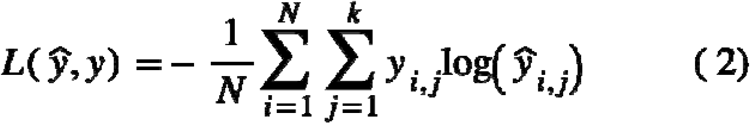

Cross entropy loss measures the number of bits needed to represent the difference between the target label and the model prediction by treating both as discrete probability distributions. Ideally, there would be no difference between prediction and label distributions, resulting in a score of 0. Prediction uncertainty, which is represented by a probability of less than 1 for the target species class of the prediction, results in higher scores. In this way, the metric measures the degree of uncertainty in the predictions of the model. Cross entropy loss is a good measure of model robustness particularly in cases where new classes are introduced into the test set because the metric can capture how the model responds to an increase in test data entropy. Models that have a stronger ability to differentiate between learned classes and new classes have lower cross entropy loss scores and can be considered more robust. Weighted F1 score is similar to the macro-F1 score. Weighted F1 score is the F1 score for each species class weighted by the support for each species class divided by the number of instances in the dataset, N. The equation for weighted F1 score is given by Equation (3). Where macro F1 score weights each species class equally, weighted F1 is based on the number of samples per class (i.e. abundances of species) in the test dataset. This means that accuracy of species that are more common in the dataset has a greater influence on the weighted F1 score than the accuracy of rare species.

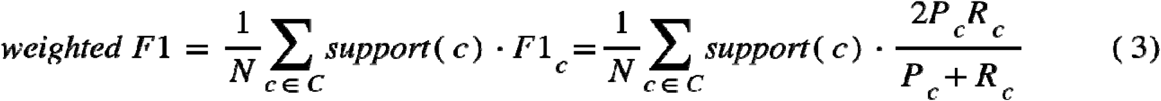

Finally, full confusion matrices were calculated for each submission to allow for further analysis, discussion and comparison, particularly to identify classes that are commonly confused (e.g., species within a genus) across methods.

Evaluation scores and confusion matrices were calculated with the scikit-learn package for python (Pedregosa et al., 2011). The evaluation code is available in the supplementary material.

### Classification algorithms

A gamut of classification algorithms were used in this competition, with the majority of teams favoring a deep learning approach (Table 2). The winning method from the previous competition, which used the HSI data in a PCA-based dimensionality reduction in a random forest gradient boosting ensemble, was used to generate baseline results (Anderson, 2018). The Fujitsu Satellite team’s method used a two-step process. First, a neural network was used to encode pixel HSI data as a 2048-dimension feature vector. The data was clustered using K means then used to create crown level feature vectors. Finally, the crown level features were put through a 3 layer fully connected neural network with RELU activation and softmax output layer for the final species classification. The Jeepers Treepers team also used a neural model, but unlike the Fujitsu Satellite method, the method fused RGB, HSI, and LiDAR data. The Jeeper’s method first used the ResNet convolutional neural network on RGB crown data to produce a species probability vector. This vector was concatenated with HSI data from the centroid of each crown and pseudo-waveform LiDAR returns. The concatenated vector was fed through a two-layer multi-layer perceptron with a customized soft-F1 loss function for final classification. Más JALApeñoS team’s method applied extreme gradient boosting to HSI data that was first filtered at a pixel level using LiDAR heights. The height-filtered pixels were further filtered using PCA based outlier removal before application of PCA based dimension reduction. The dimensionally reduced data was run through the extreme gradient boosted model and crown pixels were averaged to make the final species classification. Finally, the Intellisense CAU team’s method was based on a 1-D convolutional neural network (CNN) applied to HSI data. The CNN consisted of a convolutional layer, max-pooling layer, a fully connected layer and output. The output of the CNN was filtered using LiDAR data to remove ground pixels.

## Results

### Delineation results

Fujitsu Satellite’s multi-sensor neural network approach had the strongest performance in the delineation task with a mean IoU of 0.63 and mean RandCrowns Index of 0.86 (Table 1). The next best teams scored considerably lower in both evaluation metrics. The Faster-RCNN detector method using RGB data by the Intellisense CAU team and the region growing algorithm using a single HSI band for the baseline scored similarly in IoU (0.36 and 0.31, respectively) and the RandCrowns metric (0.61 and 0.54, respectively). The mean shift algorithm using the LiDAR point cloud by the INRAE-GIPSA team had the lowest performance (IoU=0.18, RandCrowns=0.21). The multi-sensor approach by the Fujitsu Satellite team was unique in their use of RGB, lidar, and hyperspectral data, whereas the other teams focused on only a single sensor type. The region growing baseline approach performed well relative to the two other single-sensor approaches (IoU = 0.31, RandCrowns = 0.54) even though it was run with default parameters and trained to this particular set of sites. While patterns of performance among sites were different for each method, there was no apparent trend in performance across all methods, and particularly performance was not overall lower on the TALL test site where the methods were not trained (Fig. 4).

**Table 1.** Delineation evaluation metrics for three participating teams. Metrics are the mean intersection over union (IoU) and a variation of the Rand Index that uses uncertainty halos (Stewart et al., 2021). The baseline score is from the R package, “itcSegment”, which is similar to the method used in the previous version of the competition (Marconi et al. 2019)

| Team | Method | Data used | Mean IoU | Mean RandCrowns |
| --- | --- | --- | --- | --- |
| Fujitsu Satellite | Instance segmentation pipeline | HSI, RGB, CHM | 0.63 | 0.86 |
| Intelligence CAU | Faster-RCNN detector | RGB | 0.36 | 0.61 |
| INRAE-GIPSA | Mean Shift algorithm | LiDAR point cloud | 0.18 | 0.21 |
| FEM (baseline) | Growing region algorithm | HSI - band at 812 nm | 0.31 | 0.54 |

**Figure 4.**
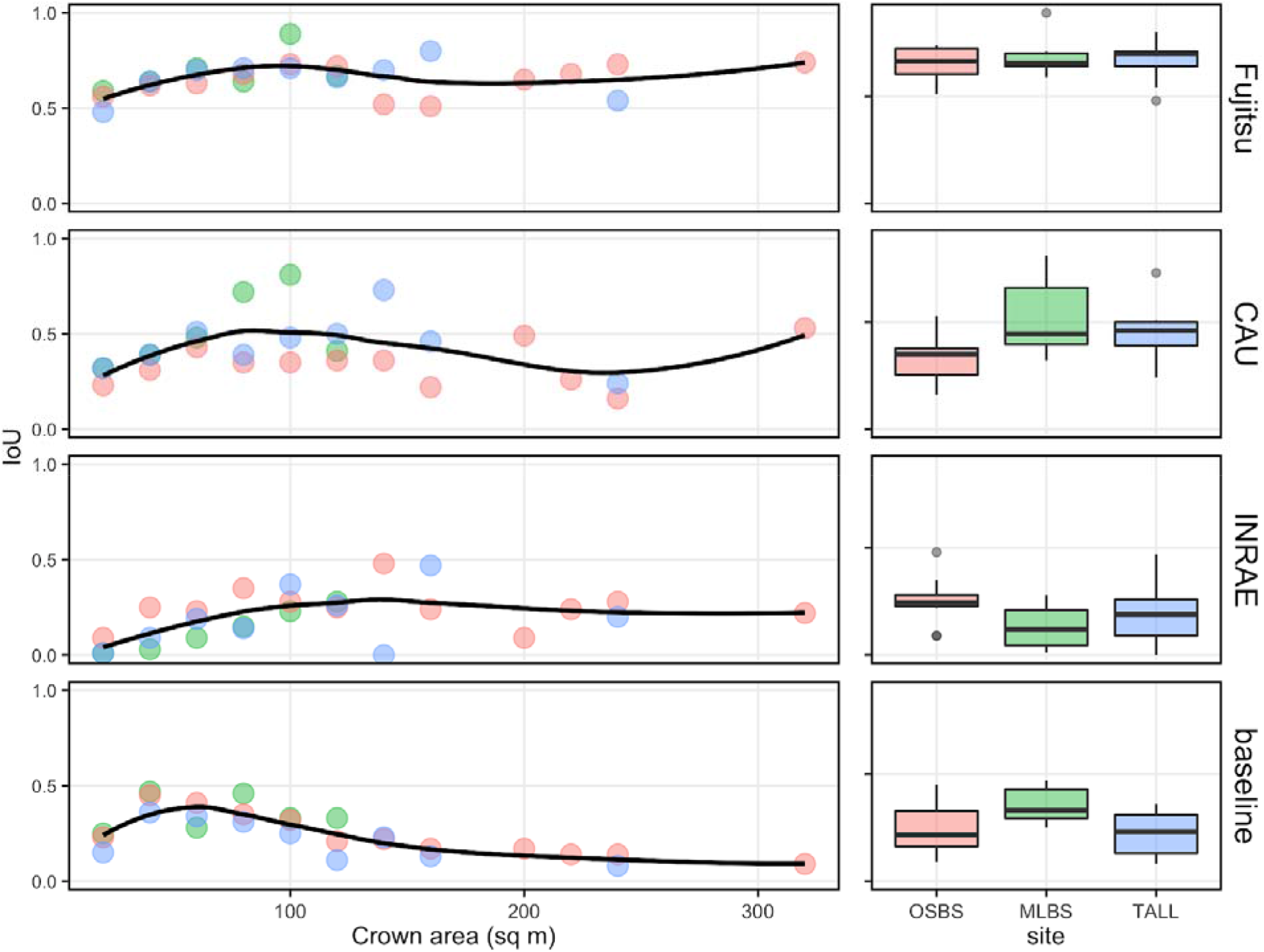
Intersection over Union (IoU) scores for each team’s delineations. Left: IoU scores vary by individual tree crown (ITC) area. Each point represents the mean IoU score for all ITCs within a 20 sq meter crown size bin, and a smoothed curve fit to all points showing the trend of scores across crown size. Points are colored by site. Right: Boxplots show differences in IoU scores across the trained (OSBS and MLBS) and untrained (TALL) sites.

Across all methods, delineation scores varied in a similar way as a function of crown size. IoU scores of individual delineations tended to be lowest for the smallest crowns, peak at crown sizes from 50 to 100 sq meters, then decline with increasing size. This pattern was especially apparent in the baseline delineations, with a peak IoU score for crowns of approximately 75 sq meters and a steady decline with increasing crown size (Fig. 5).

**Figure 5.**
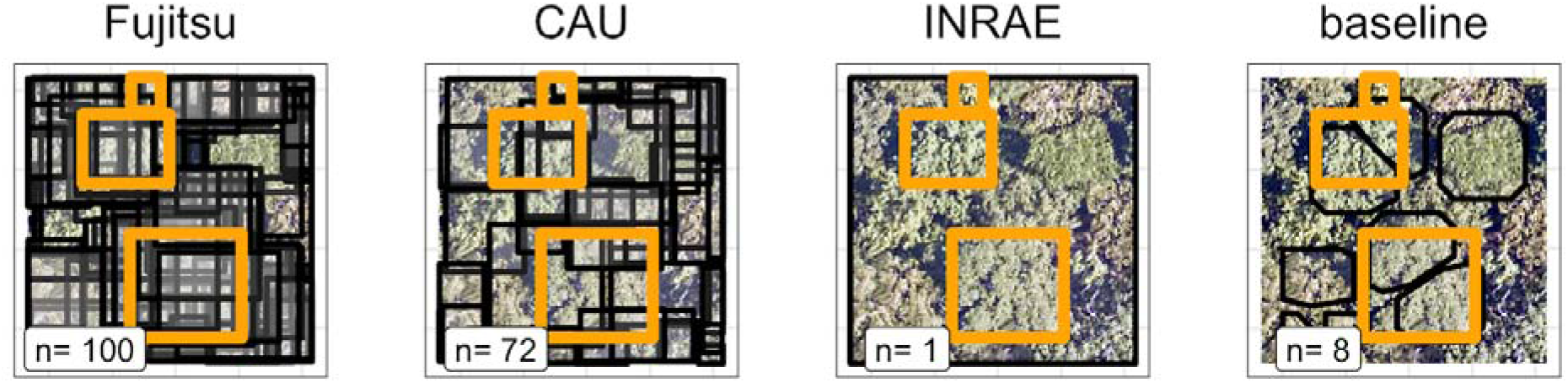
Example delineations from all teams at one plot. The “n” in the white boxes are the number of delineations for this plot for each team (shown in the black boxes, INRAE delineation is the entire plot). Orange boxes are the growth truth delineations made from visiting the site in the field and were used in the evaluation of the delineation task. The ground truth delineations are a subset of the trees in the plot.

The multi-sensor neural network approach differed from the other methods in that it produced many more delineations per plot. The average number of delineations per plot produced by the pipeline was approximately 100 for each site, whereas other approaches averaged around 30 delineations per plot with a wide range in delineations among teams. This means that the multi-sensor neural network produced far more trees than were actually present in the plot (Fig. 5), which potentially makes these predictions not suitable for many kinds of ecological analysis. This is also likely to increase the scores of this method relative to other approaches.

### Classification results

#### Performance on sites without training data

The Fujitsu Satellite team’s two-stage fully connected neural network approach had the strongest performance in the classification task for two evaluation metrics (Macro F1 = 0.32 and Weighted F1=0.53) and the second-best performance in the cross-entropy metric (Log-Loss=3.6, Table 2). The rank of models from best to worst performance was consistent for both Macro F1 (a measure of the proportion of species correctly classified) and Weighted F1 (a measure of the proportion of individual trees correctly classified), all methods performed considerably better in Weighted F1 than Macro F1, and there was greater similarity among methods in Weighted F1 than Macro F1 (Table 2). The model rank was different when evaluated with cross-entropy, which incorporates the accuracy of uncertainty for predictions, with the Más JALApeñoS extreme gradient boosting method showing the best performance (with the lowest Log-Loss lowest value of 2.4). Most methods performed better than the baseline random forest and gradient boosting ensemble method in Macro F1 and cross entropy, and all teams performed better than the baseline method in Weighted F1.

**Table 2.** Classification evaluation metrics for participating teams. Scores are from test data from only the OSBS and MLBS sites. The baseline method implemented was the Stanford-CCB approach (Anderson 2018) and was the winning method from the previous version of the competition that only used data from the OSBS site (Marconi et al. 2019). For cross entropy lower scores are better.

| Team | Method | Data used | Macro F1 | Weighted F1 | Cross entropy |
| --- | --- | --- | --- | --- | --- |
| Fujitsu Satellite | Two-stage fully connected neural network | HSI | 0.32 | 0.53 | 3.6 |
| Intelligence CAU | 1D-CNN | HSI, CHM | 0.24 | 0.50 | 7.0 |
| Más JALapeñoS | Extreme gradient boosting | HSI and canopy height for ITCs with field data | 0.14 | 0.40 | 2.4 |
| Jeepers Treepers | Two-stage neural network: RetinaNet + multimodal neural network | RGB, HSI, LiDAR point cloud | 0.09 | 0.43 | 9.2 |
| Stanford-CCB (baseline) | Random forest and Gradient boosting ensemble | HSI | 0.13 | 0.39 | 7.4 |

#### Transferability

All methods performed better on the data from the trained sites (OSBS and MLBS) than on the data from the untrained site (TALL, Fig. 6). The two-stage fully connected neural network model with the highest Weighted F1 score (Fujitsu Satellite) dropped to from 0.53 to 0.27 when the method was applied to data from only the TALL site. This lower performance on the TALL site was consistent across all methods and F1 score metrics. While Log-Loss scores were better (lower) for all teams on the trained sites, the two methods with the lowest scores performed similarly for the trained and untrained sites (Fujitsu Satellite and Más JALApeñoS).

**Figure 6.**
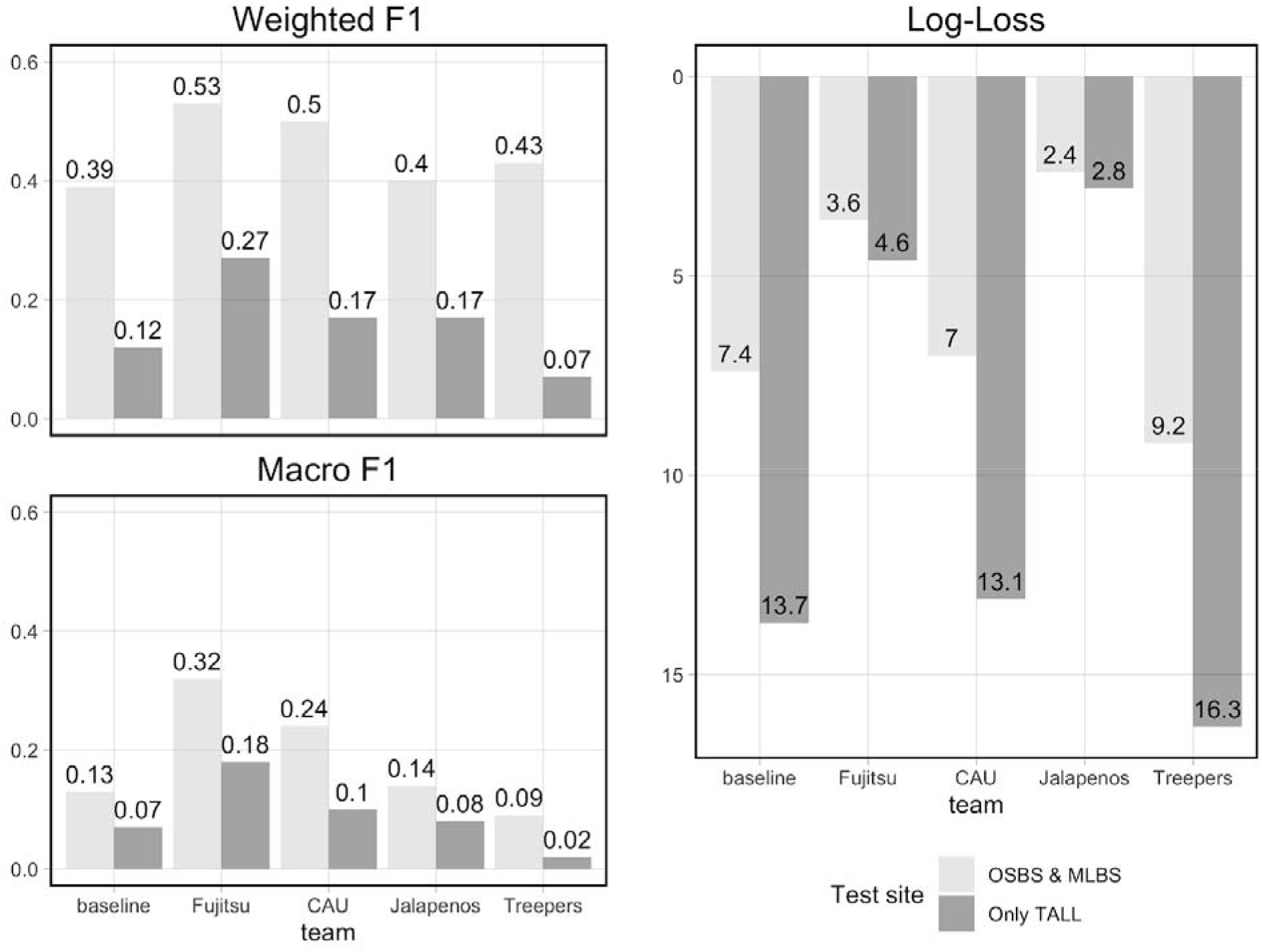
Classification evaluation metrics for all teams and different evaluation site data. Higher scores for Weighted and Macro F1 and lower scores for Log-Loss are best. The baseline method implemented was the Stanford-CCB approach (Anderson 2018) and was the winning method from the previous version of the competition that only used data from the OSBS site (Marconi et al. 2019)

#### Results by species

Model performance varied widely for predictions of individual species classes, with the general pattern of better performance for the most common species and poorer performance for the least common species (Fig. 7). Because of the high number of taxonID classes, we focus on a subset of classes that highlight important patterns seen across teams and how predictions compare for the trained sites (OSBS and MLBS) and untrained site (TALL). The most common species in the dataset (Fig. 3) is PIPA2 (*Pinus palustrus*, or Longleaf pine by its common name), which is a dominant canopy species in the conifer forests in parts of OSBS and TALL. For all methods, PIPA2 was the best-scoring taxonID, with F1 scores ranging from 0.67-0.84 across all sites. In addition, recall for PIPA2 is higher than precision for all but one team, which means that most models tend to over predict PIPA2 relative to other species (recall and precision data not shown). As with the overall accuracy metrics, all methods, including the baseline approach, predicted PIPA2 more accurately at the trained sites (F1=0.72-0.86) than at the TALL site (F1=0.12-0.54).

**Figure 7.**
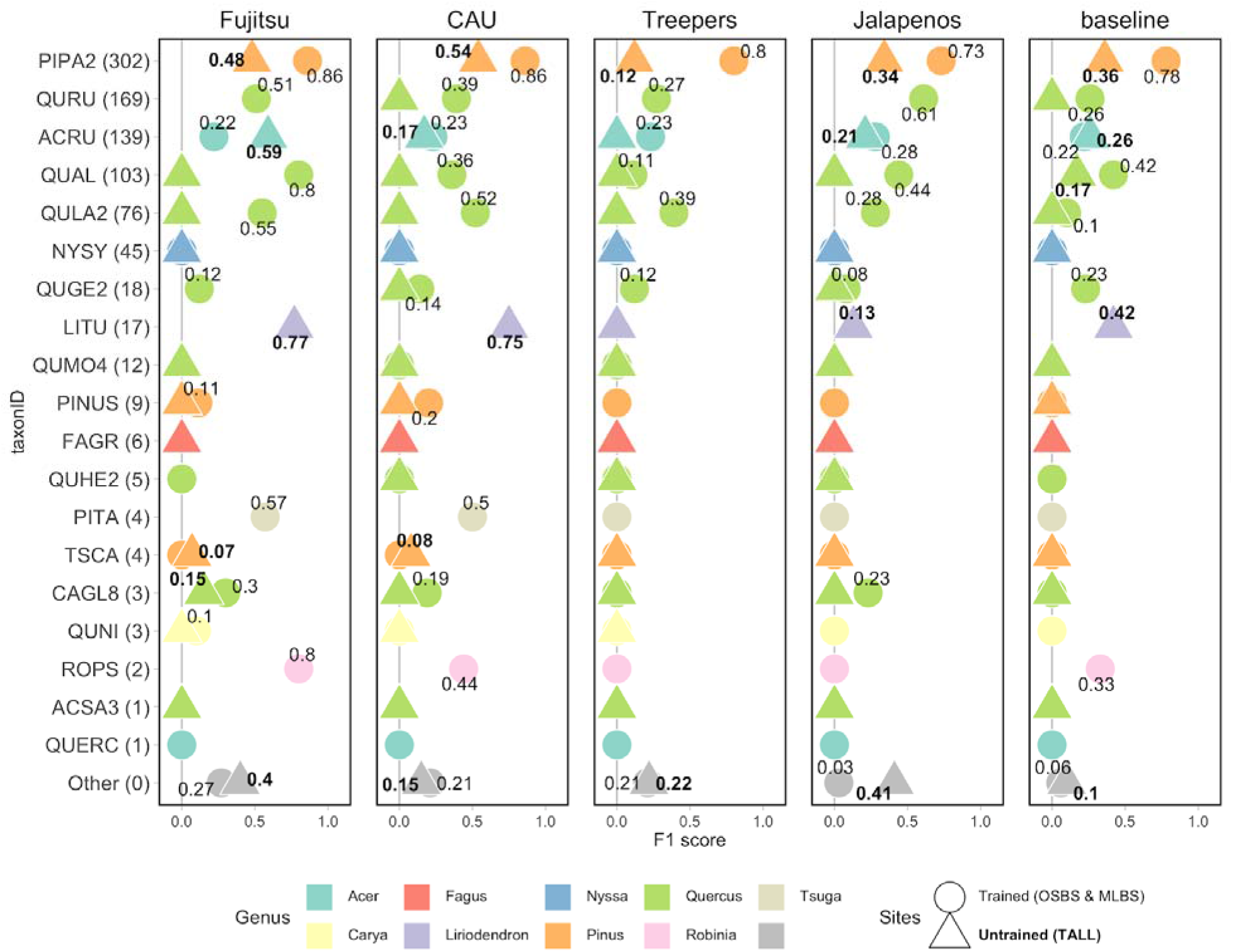
F1 score for species classes and teams. TaxonID classes are in order of highest to lowest number of samples in the training data with number of samples in parentheses. The “Other” category includes predictions of taxonID classes that were not present in the training data but were present in the testing data, and therefore represent the ability for a method to identify unknown or untrained classes. The shape represents which sites are included in the evaluation (circle=trained sites, OSBS & MLBS; triangle=untrained site, TALL). Full species information for taxonID labels are in Appendix A.

The two-stage fully-connected neural network approach (Fujitsu Satellite) had the strongest performance of the teams in predicting some of the less abundance species (Fig. 7): *Robinia pseudoacacia* (ROPS, Black locust), *Liriodendron tulipifera* (LITU, Tulip tree), *Tsuga canadensis* (TSCA, Eastern hemlock) and *Acer rubrum* (ACRU, Red maple). The strong predictions for these species, versus a focus on high prediction of dominant species, contributes to the Fujitsu Satellite team’s strong overall macro F1 score. For example, LITU has a low number of samples in the training data (Fig. 3), is taxonomically distinct as the only species in the *Liriodendron* genus, and occurred in the OSBS/MLBS training data, but was only in the test data for TALL. The strongest methods (as measured by F1; Fujitsu Satellite and Intellisense CAU) had high F1 scores for LITU at TALL in comparison to other teams and the baseline (Fig. 7), highlighting that classification approaches differ in terms of transferability of predictions to untrained sites. In addition, the Fujitsu Satellite team had 100% precision for 2 less abundant species, QUAL and TSCA, though this came at the expense of lower recall as this algorithm overpredicted the prevalence of QUAL and TSCA (precision data not shown, but see Appendix B for team confusion matrices).

In addition to comparing classification methods, the aggregated confusion matrix (Fig. 8) shows patterns of misclassification, which is important for understanding the data and how to improve models in the future. For example, inaccuracy in PIPA2 predictions was mostly due to confusion with a taxonomically and structurally similar species or with a species that co-occurs in the same habitat. The number of total correct PIPA2 predictions was 767 (66% of the total PIPA2 predictions) for all teams. For 69 predictions (5%) of the total PIPA2 predictions for all teams, PIPA2 was predicted when the true species was PITA (*Pinus taeda*, or Loblolly pine), which is a structurally similar species in the same genus. While two methods correctly predicted PITA at the TALL site, (F1 scores were very low at 0.07-0.08), all other methods and the baseline did not predict any PITA, suggesting that most methods were not sensitive to the spectral differences between these two classes (PITA and PIPA2). For 70 (7%) of the total PIPA2 predictions for all teams, PIPA2 was predicted when the true species was QULA2 (*Quercus laevis*, or Turkey Oak), which is in a different genus (oak) and is a broadleaf species (versus needleleaf of the Pines), but co-occurs in the same habitat. This species confusion seen by many methods could be due to similarities in the spectral features identified by the models, by errors in labels of pixels due to overlapping tree crowns, or by similarities in background reflectance influencing the predictions.

**Figure 8.**
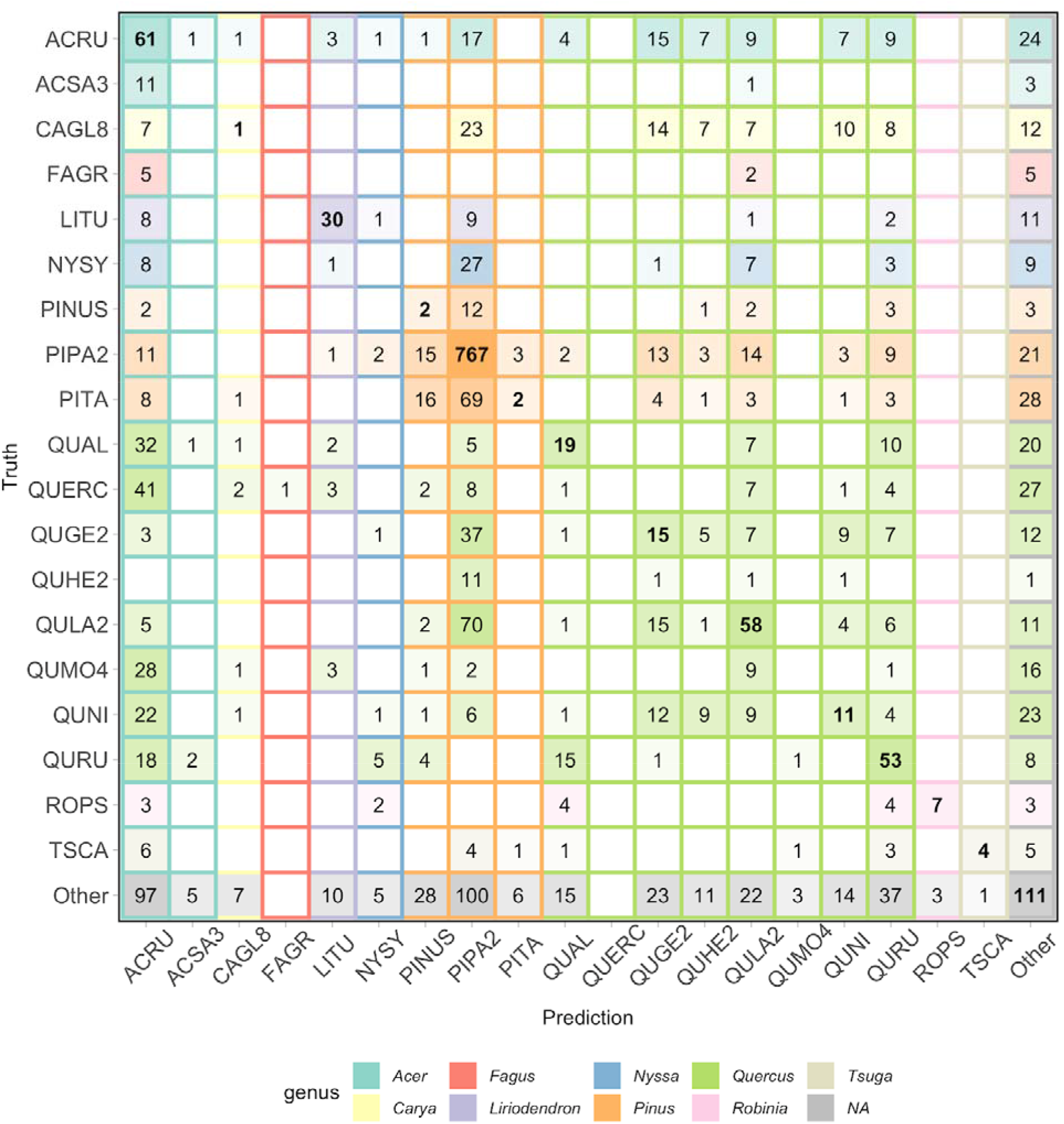
Aggregated confusion matrix of all team predictions. Numbers with the cells and intensity of color represent the total number of predictions for a given ground truth label. Correct predictions are along the diagonal with bold text. Background color corresponds to the true genus category and box outline corresponds to the predicted genus category. Confusion matrices by team are in Appendix B.

When making predictions for the “Other” class, all methods performed better than the baseline in their ability to detect untrained species classes. Three teams performed equally in predicting the “Other” species class at the OSBS and MLBS sites with F1 scores of 0.21-0.27 (Fig. 7). The baseline method and the JALApeñoS team performed poorly in identifying new species with F1 scores of 0.06 and 0.03. An encouraging result is that two methods (Fujitsu Satellite and MaS JALApeñoS) had high F1 scores of 0.40 for the “Other” species class at the TALL site (Fig. 7), highlighting that these methods are able to identify new classes when applied to a new site.

## Discussion

### Delineation

This competition compared four approaches for delineation of individual tree crowns that produce widely differing results. Two teams developed methods that showed promise for advancing delineation methods. The team with the highest scores (Fujitsu Satellite) used a complex instance segmentation pipeline that required the integration of three types of remote sensing data. The number of delineations was not appropriate for use in ecological studies because there were multiple possible delineations of a single tree crown, resulting in far greater numbers of trees than were present in a plot. Therefore, an instance segmentation pipeline shows promise if coupled with a method to select the best delineation from the sets of overlapping delineations for each tree. An example is the recent work on a DeepForest method that generates multiple delineations, and then uses a non-max suppression to output a single final delineation for each tree (Weinstein et al., 2020a). A limitation of operationalizing the instance segmentation pipeline approach is the reliance on three remote sensing data types (HSI, RGB, and CHM). While these datasets are available for the NEON sites, this is not typical for many areas. The second highest score was earned by the Intellisense CAU team that used only RGB data with a Faster-RCNN detector approach. While the RCNN approach is complex, only having the RGB data requirement is encouraging because RBG data is much more widely available than LiDAR and hyperspectral data, potentially allowing RGB based methods to be applied more broadly (Weinstein et al., 2020b). Finally, the mean shift algorithm on the LiDAR point cloud did not perform as strongly as the other approaches for two main reasons; low point density and the influence of forest composition. First, the mean shift algorithm benefits from high point density (20 and 60 pulses per m2, Tusa et al. 2020) providing abundant information on the spatial distribution of the neighbors of each point (Wu, Yao & Polewski, 2018). A low point density does not provide enough definition among clusters, which leads to undersegmentation, as was seen in the results. An approach to overcome the challenge of low point density is the fusion of the spatial information from a LiDAR point cloud with the RGB image of high spectral and spatial resolution by introducing additional kernels (Comaniciu & Meer, 2002). Second, the kernel shape implemented in the algorithm is a superellipsoid, whose profile is more suitable for delineating conifers rather than hardwood trees. Although proper parameter setting can improve tree delineation, prior information concerning the spatial distribution of the tree species is critical to adapt the kernel profile shape and size.

The results from this competition, as well as recent methods comparisons (Aubry-Kientz et al., 2019), highlight the need for continued work in tree crown delineation from remotely sensed imagery. A key component will be the availability of open-source models and benchmark evaluation data to compare future methods on a standard set of data. A tree crown detection benchmark was recently published which will enable future users to compare methods across multiple NEON sites (Weinstein et al., 2021). This dataset, combined with the deep learning model made available in the DeepForest package (Weinstein et al., 2020a), can serve as a baseline for future methods development.

One lesson learned from this competition is the importance in selecting an appropriate evaluation metric to accurately reflect the strength of delineations that produce valuable outputs for ecological studies. For each ground ITC, our evaluation pipeline selected the best of all predicted delineations and used this best delineation to calculate two evaluation metrics (IoU and RandCrowns). As a result, our evaluation favored methods with multiple candidate predictions for each crown since there was no penalty for multiple predicted delineations for each crown. This evaluation pipeline may have increased the apparent strength of the winning method. While we do believe that the instance segmentation pipeline by the Fujitsu Satellite team is promising, especially coupled with non-max suppression, the results illustrate that evaluation metrics should be chosen carefully to focus on outputs relevant to the application, in this case mapping canopy trees for ecological analysis. Standard metrics such as IoU can be used, but evaluation pipelines should penalize for too many overlapping delineations, such as scaling the IoU or RandCrowns by the number of candidate detections, selecting the worst rather than the best intersecting detection, or penalizing the score for each “extra” detection. This may be advantageous when evaluating applications where detecting the approximate location and size is more important than accurate delineation for measuring crown size parameters. In addition to evaluation metrics that assess delineation accuracy of individual crowns, evaluation metrics such as detection rate that assess the accuracy of the number of ITC delineations on a plot level could be used to reward methods produce the correct number of delineations over larger areas as well as assess metrics of interest to ecologists and foresters (Yin & Wang, 2016).

A challenge worth noting is that the training and testing data were generated in two different ways. The training data was generated by the IDTReeS research team using multiple remote sensing images and NEON field inventory data as support. Every apparent individual tree crown in the plots was delineated, even those that were uncertain due to their small size or adjacency to other crowns. This introduced some uncertainty in the training data because the delineations were not generated or validated in the field. The test data, in contrast, were crown delineations that were generated in the field and only crowns that were clearly identifiable in the remote sensing imagery were delineated. Therefore, the test data had greater certainty and did not include very small crowns or those that were not apparent in the imagery. In this way, the performance of the methods in this competition may be inflated because operational tree crown delineation requires all crowns, even small crowns or crowns that are not fully in the canopy, require delineations.

### Classification

Most remote sensing approaches to tree species classification focus on a method for a single site and where all species classes are known. In this iteration of the competition, we asked participants to grapple with more challenging classification tasks, specifically building models using training data from multiple sites and applying those models to a novel location. Our results show the promise for all methods to generalize by learning from data from multiple sites, but all methods had a limited ability to produce strong predictions for the untrained site. Two classification methods using neural networks stood out due to their ability to predict common classes and learn unique spectral features of uncommon species classes. The first-ranked team based on F1 scores (Fujitsu Satellite) used a two-stage fully-connected neural network approach, and the second-ranked team (Intellisense CAU) used a 1-D convolutional neural network. These methods also performed well based on the Log-Loss score, but the first-ranked team based on Log-Loss (Más JALApeñoS) used an extreme gradient boosting approach.

The classification task for this competition was challenging due to limitations and complexities of the data; complexities that reflect the characteristics of data for real-world applications for which robust methods are needed. One common challenge for ecological applications is that the amount of field data for training and testing is often smaller than the optimal amount to train and robustly evaluate algorithms. We believe the most accurate data for training and evaluating crown delineation and classification models comes from laborious field efforts where individual tree crowns are delineated and species are identified in the field (Graves et al., 2018). Datasets like these are small and often limited to specific sites and studies. To overcome limited field data and create a sufficiently large dataset for this competition, we generated a large set of image delineated crowns to use as training data (see Appendix A). The certainty of these image delineated ITCs, especially for classification, is less than for the field data because of uncertainty in spatially aligning field data of individual trees to the remote sensing data. The results showing confusion between two very different species (Fig. 8) suggest that some of the training pixels identified as belonging to one species may in fact belong to another, presenting challenges to classification models. Therefore, the extra challenge of this competition is learning from data with uncertain labels. We emphasize that this is an inherent challenge in ecological studies since high-quality data, such as the field-delineated ITCs, will always be limited, and therefore there is a need for methods to account for this source of potential uncertainty. Future efforts should be made to support improved alignment between field and remote sensing datasets. In generating datasets, field data could indicate if a tree has a position in the canopy and is therefore viewable in remote sensing imagery. Additionally, tree crowns could be digitized in remote sensing data while in the field to avoid any uncertainty and build robust datasets (Graves et al., 2018). Future efforts in developing more robust models to handle data with potential misalignment, the active research in image analysis in classification and detection with label uncertainty (Zou, Gader & Zare, 2019; Du & Zare, 2019) may provide a way forward for these tasks.

Another inherent challenge with the classification of ecological data is the imbalance in the data across classes. Most natural forest ecosystems have an unequal abundance distribution of species, with a “hollow curve” shaped distribution where there are a small number of common species, and a large number of relatively rare species (McGill et al., 2007). Ecological datasets often reflect this natural distribution since they are generated by randomly sampling plots in the field. Our results reflect the common outcome of classification on unbalanced datasets, with models generally performing better on classes that have a greater representation in the training data compared to classes with lower representation. Evaluation metrics that are weighted by the number of samples per class, such as Weighted F1 which was one metric used in this competition, favor models that are most accurate for abundant classes. However, for many ecological questions and applications, having strong predictions across all species, especially the rare species is important (Leitão et al., 2016; Dee et al., 2019; Cerrejón et al., 2021), and therefore an evaluation score such as Macro F1 is most appropriate. Understanding patterns of taxonomic and functional diversity or evaluating the impact of climate changes and extreme disturbance events on species are examples where poor accuracy of rare species will impact the ability to use the predictions because of the uncertainty in the predictions. This imbalance of the data impacted all teams, but methods using neural networks show promise for predicting less abundant species (Fujitsu Satellite and Intellisense CAU, Fig. 4).

Finally, transferability to new sites is challenging because the new site will most likely contain new taxonomic classes and introduce spectral variation, especially if the new site is geographically distinct from the training sites. We found that across metrics (F1 and Log-Loss), all methods performed worse on the untrained than the trained sites. While the decreased performance is partially due to the change in species classes, even a dominant species (Longleaf pine, PIPA2) was more poorly predicted at the untrained site. This suggests that regardless of differences in species presence and abundance between sites, the spectral and structural signatures of individual species (caused by sensor calibration, atmospheric conditions, seasonal differences, or inherent differences in species foliar and structural properties) are sufficiently different to hinder model performance. An encouraging result was the presence of methods with significantly lower Log-Loss scores (gradient boosting by Más JALApeñoS and a neural network by Fujitsu), and that scores were similarly low for both the trained and untrained sites (Fig. 6). A low Log-Loss score means that a method was confident with its correct predictions and unconfident with its incorrect predictions. Methods that score low in Log-Loss could be most useful in transferring to new sites because low confidence in a prediction could indicate the presence of a new species.

Another way we assessed transferability was by measuring each methods’ ability to identify new classes at both the training sites as well as the novel test site by having teams predict an “Other” species class. While the weighted F1 scores for the “Other” species class are too low for accurately detecting these new species (∼0.2, Fig. 4), most methods performed better at this task than our baseline approach from the previous competition where transferability was not part of the task. In addition, methods generally performed better at identifying new species in the trained sites (OSBS and MLBS) than the untrained site (TALL). The ability for methods to detect new species is encouraging because it shows an advancement of methods, especially since the previous competition in 2017, to be able to handle the operational scenario where species exist on the ground that are not part of the training data.

## Conclusions

This competition challenged the data science, remote sensing and ecology communities to train and apply algorithms for the common tasks of delineation and classification in the context of individual tree detection and species identification across multiple sites. A wide range of delineation methods were implemented in this competition, showing you can approach this task in multiple ways and have variable results. There is great potential for improvement of delineation methods, especially for small and large crowns, but methods generally work equally well regardless of the sites they are trained and then applied. Participants also developed a wide range of approaches to classification of tree crowns to their taxonomic species, many of which were significantly better at cross-site prediction than the best method from previous competition. Most methods can predict common classes well, even across sites, but more work is needed in methods that can handle imbalanced data and learn from rare species, i.e. those with lower relative abundances, and are robust to new species when applied to an untrained site. This competition has highlighted the value in comparing methods on a standardized dataset to implement approaches from a broad range of expertise, and highlight areas where we can continue to improve.

## Acknowledgements

Thanks to Morteza Shahriari Nia for helping create the initial IDTReeS competition. The National Ecological Observatory Network is a program sponsored by the National Science Foundation and operated under cooperative agreement by Battelle. This material is based in part upon work supported by the National Science Foundation through the NEON Program. This research was supported by the Gordon and Betty Moore Foundation’s Data-Driven Discovery Initiative through grant GBMF4563 to E.P. White, by the USDA National Institute of Food and Agriculture McIntire Stennis project 1007080 to S.A. Bohlman, and by the National Science Foundation through grant 1926542 to E.P. White, S.A. Bohlman, A. Zare, D.Z. Wang, and A. Singh and grant 1442280 to S. A. Bohlman.

## Notes

https://zenodo.org/record/3934932

https://github.com/sjgraves/IDTReeS_competition

https://github.com/weecology/idtrees_competition_evaluation

### Competing Interest Statement

The authors have declared no competing interest.

