## Appendix A for "Data science competition for cross-site delineation and classification of individual trees from airborne remote sensing data"

#### Details of ITC and field data generation

This document provides additional information about the Individual Tree Crown (ITC) delineations and field data provided with the IDTReeS Competition.

### NEON data products

**Table A1. Data products from the National Ecological Observatory Network (NEON) used in the competition.**

| Data product | Description | Spatial resolution | Data format provided to participants |
|---|---|---|---|
| High-resolution orthorectified camera imagery (RGB) | Raster data of the reflected energy from the surface as 3 bands representing the red, green, and blue portions of the spectrum. NEON data product ID: DP1.30010.001 | 10 $cm^2$ | GeoTiff (.tiff) |
| Discrete return LiDAR point cloud (LAS) | Point cloud data of values in the X,Y, Z direction of the height of surface features and the ground. NEON data product ID: DP1.30003.001 | 6 points per $m^2$ | .las |
| LiDAR canopy height model (CHM) | Raster data containing the height of the top of the vegetation canopy. The CHM is a product of the LiDAR point cloud data. NEON data product ID: DP3.30015.001 | 1 $m^2$ | GeoTiff (.tiff) |
| Mosaic hyperspectral surface reflectance (HSI) | Raster data of reflected energy from the surface as 426 5-nm wide wavelength bands from 380-2510 nm. NEON data product ID: DP3.30006.001 | 1 $m^2$ | GeoTiff (.tiff) |
| Field measurements | Attributes of individuals measured in the field as part of the Woody Plant Vegetation Structure sampling protocol. NEON data product ID: DP1.10098.001. | Individual tree | .csv |
| Individual tree crown (ITC) boundaries | 2-dimensional rectangular bounding boxes that geographically defines an ITC. Data are not standard NEON data products and were generated the IDTReeS research group. | Individual tree | Esri Shapefile or .csv file with WKT format |

### Individual Tree Crown (ITC) data

ITC data are fundamental for both tasks in the competition. For delineation participants are given labeled ITCs in the training dataset and generate ITCs for the test dataset. For classification, participants are given ITCs labeled with the taxonomic species labels, and generate taxonomic species predictions for unclassified ITCs.

ITC data were generated by the IDTReeS research team and are not standard data products provided by the National Ecological Observation Network (NEON). The ITC data were generated with two different approaches and are used in distinct ways in the competition. Because ITC data are time-intensive and difficult to generate, the research team used a combination of two different approaches to produce both a reasonably large number of labeled data points for training and precise data points for testing. Both ITC datasets were generated by experts who are familiar with the ecology of the sites.

**Field ITCs** were generated by members of the research team by visiting each NEON site and directly mapping ITCs in the RS data while in the field. Remote sensing (RS) data was loaded onto tablet computers that were equipped with GPS receivers. While in the field, researchers digitized crown boundaries based on the location, size, and shape of the crown seen in the field onto the RS data. Complete information for how the field ITC polygon data were generated are documented in Graves et al. 2018 (https://peerj.com/preprints/27182/).

Field ITCs were originally delineated as polygons to precisely match the irregular shape of each crown. For this competition, these polygons were converted to bounding boxes to capture the maximum width of the crowns in the North/South and East/West directions (Fig. A1). Field ITCs were collected in 2015 on remote sensing data from 2014, or collected in 2016 on 2015 data. The field ITCs were verified in the 2018 and 2019 imagery that are used in this competition. Any differences between the ITCs and the most recent imagery were corrected by shifting and or adjusting the dimensions of the ITC to match the position of the crown in the imagery. If differences could not be resolved, ITCs were removed from the dataset.

The field-ITCs are considered the most accurate validation data and are used as the test and evaluation data for the delineation and classification tasks. For the classification task, species labels were generated by identification in the field or by collecting a voucher sample to be identified in an herbarium by a trained botanist. For individuals that could not be identified to species in the field or where a voucher could not be collected, the individual was identified to its genus category.

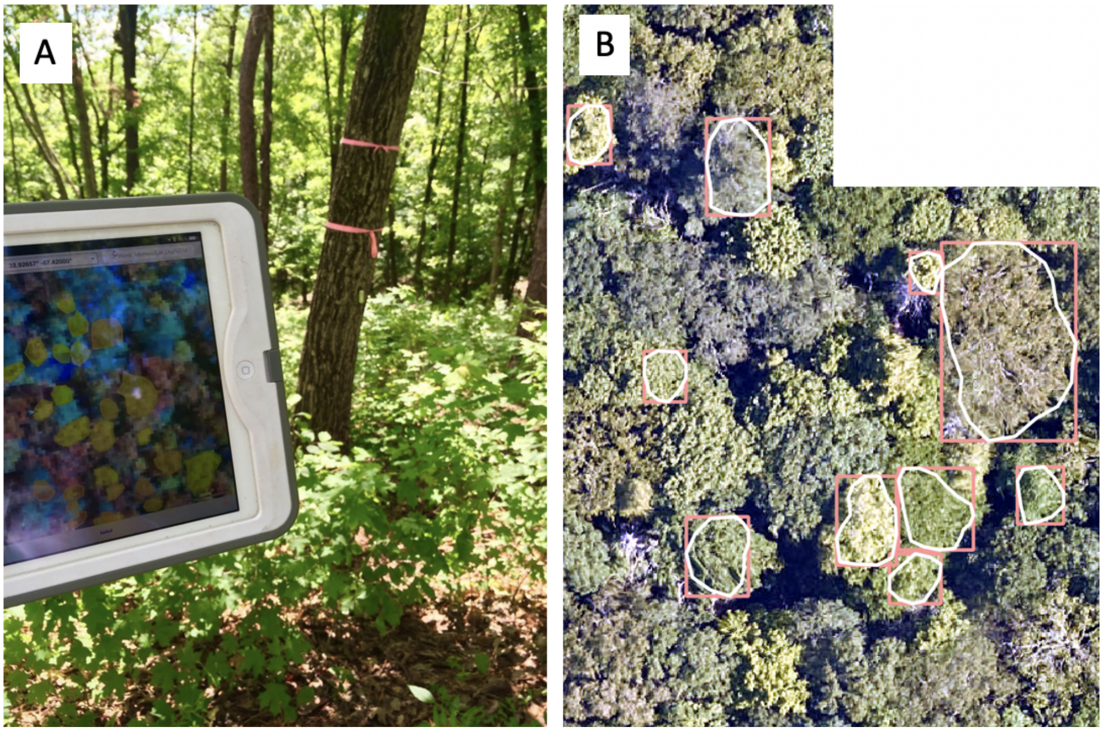

**Figure A1.** Process and examples for generating field Individual Tree Crowns (ITCs). A) Tablet computer used for digitizing tree crowns in remote sensing data while in the field. B) Examples of ITCs on 2019 high-resolution RGB camera remote sensing data. Crown polygons (white) were mapped in the field and converted to bounding box extents (pink) to use as test data for the competition. Horizontal distance in this image spans 40 meters.

**Image-only ITCs** were generated by members of the research team by identifying tree crown boundaries using all available NEON RS data. These ITCs were based on viewing lidar, RGB and hyperspectral images overlaid with point locations of stems and their attributes from NEON and using expert knowledge to visually estimate the boundaries of individual tree crowns (Figure S2). The field sites were not visited to generate image-only ITCs.

The image-only ITCs are considered by the IDTReeS research team to be the best available data that can be generated without observing individual trees in the field. Image-only ITCs are used as training data for the delineation task, and define the crown boundaries for the classification task. For the classification task, the species labels were generated by combining the ITC bounding box with NEON vegetation structure data (see next section). Using stem locations, crown sizes, and canopy position data from the NEON vegetation structure data, tree stems were matched to bounding boxes to provide species labels to each bounding box. Since all boxes needed to be assigned with an identifier, crowns not corresponding to any individual tree in the NEON vegetation structure were assigned with an arbitrary unique ID. This means that some ITCs do not have attribute data in the field data table.

Complete information for how the image-only ITCs were generated and assessment of the accuracy among expert digitizers is documented in Weinstein et al. (2020).

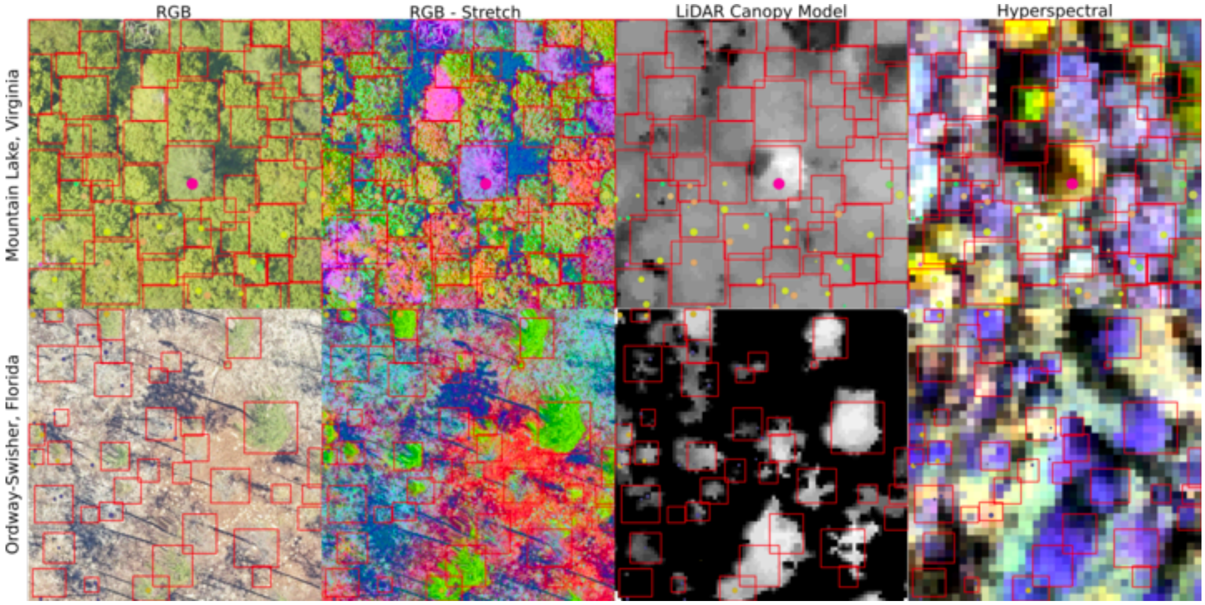


**Figure A2.** Example of image-only ITC delineations (red boxes) on various remote sensing products for 2 NEON sites. These remote sensing products were used to aid in the manual delineation of ITCs. Full details for how the image-only ITC delineations were created is in Weinstein et al. (2020).

**Table A2**. Summary of the two types of ITC data used in the competition

| ITC type | field | image-only |
|---|---|---|

| | | |
|---|---|---|
| How they were generated | Manual delineation of crowns in the field directly on RS data. Species information from field or voucher identification. | Manual delineation of crowns using the full set of remote sensing data, but with no field visit. Species information from NEON vegetation structure data |
| How they are used in the competition | Delineation: test data<br>Classification: test data | Delineation: train data<br>Classification: train data |
| Association with NEON plots located within NEON sites | Are not located in NEON sampling plots. | Are located within NEON sampling plots and directly associated with NEON vegetation structure data. |

### NEON field data

Field data are collected by the NEON Terrestrial Observation System (TOS) personnel. The data are collected in permanent plots that are repeatedly sampled. There are two types of plots associated with different field sampling objectives within NEON; the distributed plots that are spatially distributed throughout the NEON site and stratified by land cover type to capture the diversity of ecosystem types within each site; the tower plots, that are located within the airshed of the flux-tower. Data from both types of plots are used.

The field data used in this competition are from the Woody Plant Vegetation Structure sampling protocol (NEON 2020, protocol DP1.10098.001). The vegetation structure data was compiled using the NEONderive repository developed by members of the IDTReeS research team (https://github.com/MarconiS/NEONderive). The tabular data are provided as a comma separated value (csv) files that contains information on individual tree identifiers, location of trees relative to sampling locations (i.e. distance and azimuth from a central location), species and genus labels, and measures of relevant structural attributes (Table A3). The field attribute that is directly used in this competition is the taxonomic species information, described by the scientific name and the taxonomic identification code (Table A4).

**Table A3.** Field data to be used in the data science evaluation. The data provided are a subset of the full set of standard data collection by the NEON Terrestrial Observatory System. These attribute names and descriptions are also provided with the data.

| fieldName | description |
|---|---|
| indvdID | Domain-level unique identifier for an individual: NEON.MOD.D##.######. |
| siteID | NEON site code. Site codes used in this dataset are MLBS, OSBS, and TALL. |
| taxonID | Species code, based on one or more sources |

| | |
|---|---|
| scientificName | Scientific name, associated with the taxonID. This is the name of the lowest level taxonomic rank that can be determined |
| taxonRank | The lowest level taxonomic rank that can be determined for the individual or specimen |
| utmZone | UTM zone |
| nlcdClass | National Land Cover Database Vegetation Type Name |
| elevation | Elevation (in meters) above sea level |
| growthForm | The growth form classification: single-bole tree, multi-bole tree |
| plantStatus | Physical status of individual: live, dead, lost |
| stemDiameter | Cross-sectional stem diameter |
| height | Highest point of an individual or average height of a patch |
| maxCrownDiameter | Maximum crown diameter of the individual or patch |
| ninetyCrownDiameter | Crown diameter perpendicular to maxDiameter |
| canopyPosition | Vertical status of an individual relative to its neighbors |

**Table A4. Taxonomic species information for species classes used in the classification task.** The taxonomic species information is described by its scientific name, which includes a genus and species classification. To simplify the taxonomic species information, each scientific name was simplified to its unique taxonomic identification code.

| Taxonomic identification code | Scientific name |
|---|---|
| ACPE | *Acer pensylvanicum L.* |
| ACRU | *Acer rubrum L.* |

| | |
|---|---|
| ACSA3 | *Acer saccharum Marshall* |
| AMLA | *Amelanchier laevis Wiegand* |
| BETUL | *Betula sp.* |
| CAGL8 | *Carya glabra (Mill.) Sweet* |
| CATO6 | *Carya tomentosa (Lam.) Nutt.* |
| FAGR | *Fagus grandifolia Ehrh.* |
| GOLA | *Gordonia lasianthus (L.) Ellis* |
| LITU | *Liriodendron tulipifera L.* |
| LYLU3 | *Lyonia lucida (Lam.) K. Koch* |
| MAGNO | *Magnolia sp.* |
| NYBI | *Nyssa biflora Walter* |
| NYSY | *Nyssa sylvatica Marshall* |
| OXYDE | *Oxydendrum sp.* |
| PEPA37 | *Persea palustris (Raf.) Sarg.* |
| PIEL | *Pinus elliottii Engelm.* |
| PIPA2 | *Pinus palustris Mill.* |
| PINUS | *Pinus sp.* |
| PITA | *Pinus taeda L.* |
| PRSE2 | *Prunus serotina Ehrh.* |
| QUAL | *Quercus alba L.* |

| | |
|---|---|
| QUCO2 | *Quercus coccinea* |
| QUGE2 | *Quercus geminata Small* |
| QUHE2 | *Quercus hemisphaerica W. Bartram ex Willd.* |
| QULA2 | *Quercus laevis Walter* |
| QULA3 | *Quercus laurifolia Michx.* |
| QUMO4 | *Quercus montana Willd.* |
| QUNI | *Quercus nigra L.* |
| QURU | *Quercus rubra L.* |
| QUERC | *Quercus sp.* |
| ROPS | *Robinia pseudoacacia L.* |
| TSCA | *Tsuga canadensis (L.) Carriere* |

Data science competition for cross-site delineation and classification of individual trees from airborne remote sensing data

NEON (National Ecological Observatory Network). Spectrometer orthorectified surface directional reflectance - mosaic (DP3.30006.001). https://data.neonscience.org (accessed March 4, 2020)

Weinstein, B., Graves, S., Marconi, S., Singh, A., Zare, A., Stewart, D., Bohlman, S. and White, E.P., 2020. A benchmark dataset for individual tree crown delineation in co-registered airborne RGB, LiDAR and hyperspectral imagery from the National Ecological Observation Network. bioRxiv.
