## Appendix B for "Data science competition for cross-site delineation and classification of individual trees from airborne remote sensing data"

#### Team methods and results

This document describes the methods used by participating teams for the classification and delineation tasks. It also includes confusion matrices for the classification methods for each team.

**Table B1. List of teams that participated in the competition**. Teams participated in the delineation task (D), classification task (C), or both. Teams for the baseline methods were the winning teams from the previous competition (Marconi et al. 2019). The baseline teams did not participate in this competition but their methods were implemented on the data from this competition to compare their performance.

| Team name | Institution | Country | Task | Paper reference |
|---|---|---|---|---|
| Fujitsu Satellite | Fujitsu Laboratories | Japan | D,C | - |
| GatorSense | University of Florida | USA | C | Zou et al., 2019 |
| INRAE-GIPSA | Univ. Grenoble Alpes | France | D | Tusa et al. 2020 |
| Intellisence_CAU | China Agricultural University | China | D,C | - |
| Jeepers Treepers | CU Boulder | USA | C | Scholl et al. 2021 |
| Más JALApeñoS | Yale University | USA | C | - |
| Stanford-CCB | - | USA | C (baseline) | Anderson, 2018 |
| Dalponte | - | Italy | D (baseline) | Dalponte et al., 2019 |

### Fujitsu Satellite team

**Delineation**

We propose an instance segmentation pipeline for delineating the boundaries of ITCs on hyperspectral and RGB data. We employ state-of-the-art detection and instance segmentation models, such as Cascade R-CNN (Cai and Vasconcelos 2018), HTC (Chen et al. 2019) and DetectoRS (Qiao et al. 2020) to detect and segment the tree crowns. Firstly, we apply the hyperspectral super-resolution method (HSR, Fu et al. 2019) to obtain the same spatial resolution as the RGB image (0.1 m). HSR can leverage the texture information from RGB and the spectral information from HSI. For each image, we reduced the HSR hyperspectral cube (369 bands) into a false color RGB image (3 bands) to be used for performing tree crown detection and segmentation. These 3 bands are selected progressively with the best performance on a well-trained Cascade R-CNN model. Secondly, we design a two-branches backbone structure to utilize both the RGB and synthesized hyperspectral images, the weights of which are pre-trained with RGB images (Fig. B1). The backbone can extract image features from both RGB images and synthesized hyperspectral images to detect and segment the individual tree crowns. The two-branches are distributed symmetrically. Thirdly, we fuse the middle level features in each layer from the two branches and then transfer the integrated features to a feature pyramid network (FPN, Lin et al. 2017) module and a regional proposal network module (Ren et al. 2016), tree crown box regression and mask segmentation modules. Finally, we use the canopy height model to filter false positive predictions whose height is smaller than a threshold of 2 meters.

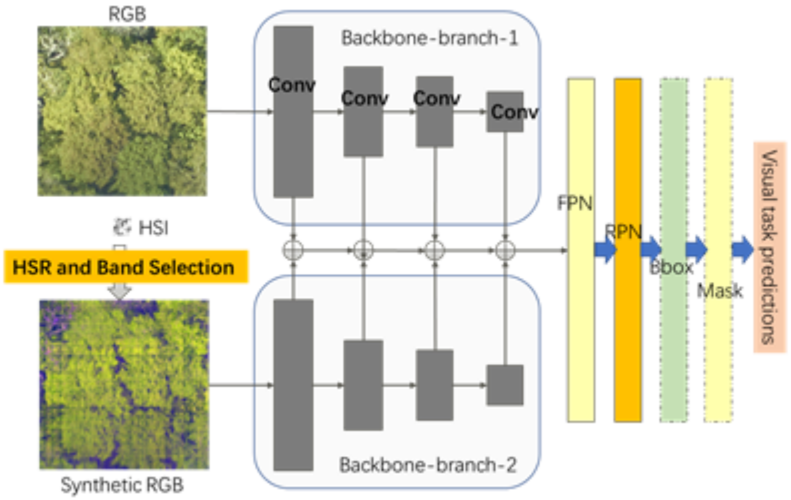


**Figure B1. Two-branches backbone structure neural network for IDTReeS detection and segmentation**

**Classification**

We propose a point-level feature-based neural network to classify the tree species using hyperspectral data. Firstly, the hyperspectral pixels are normalized by a min-max scaling method and fed into a pixel-level neural network to extract the pixel features. Figure B2 shows the structure of the pixel-level neural network classifier, which consists of four full connection layers. Rectified Linear Unit (ReLU) activation is used after the fully connected layer except the last one layer. The output dimensions of full connection layers are 2048, 4096, 2048, and the number of tree species, respectively. The third full connection layer is used as the pixel-level feature. Mixup (Verma et al. 2019) and random spatial data augmentation are used for neural network training. For the training dataset, we extract all the pixel-level features and cluster the extracted features into L clusters, set as 100 in the experiment, by K-means method. The clusters are then encoded into indexes ranging from 0 to L-1. Each index is related to a clustered feature codebook. With the L-dimension codebook, we can get L-dimension crown-level features for all crowns. The crown-level feature represents the geometry distribution of hyperspectral texture for one individual tree, which is robust for the outliers. Finally, we train a crown-level neural network to classify the individual tree crowns. The crown-level classifier is composed of two fully connected layers, the output number of which are 2048 and the number of tree species. ReLU activation is used after each fully connected layer. We can calculate the probability of each sample after the softmax layer.

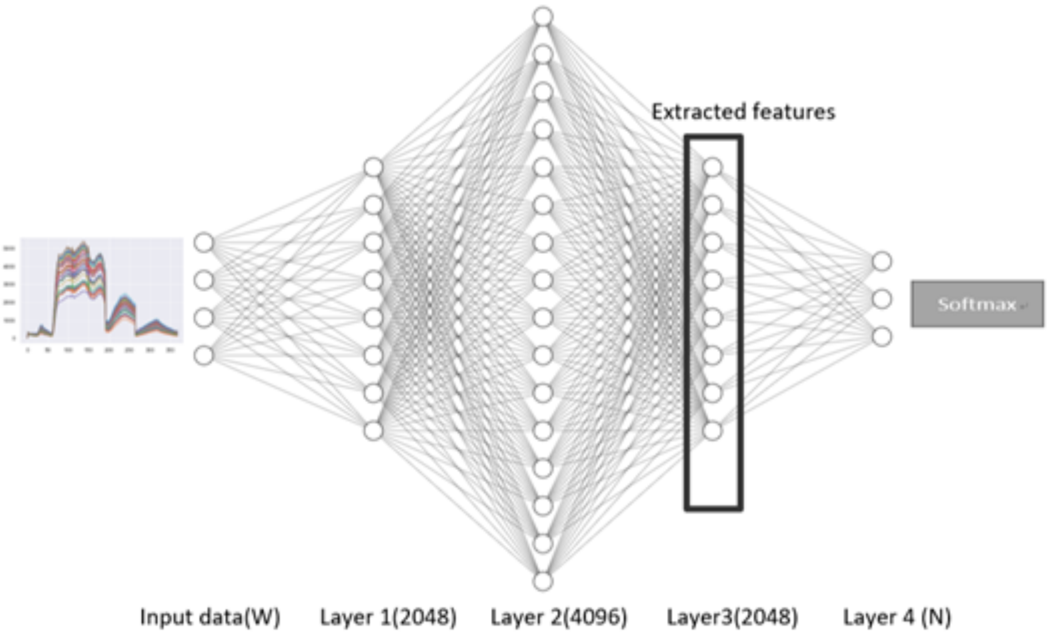


**Figure B2. Pixel-level neural network classifier.** Input is the hyperspectral of each point in the crown. Output is the tree species. The network has three fully connected layers with node numbers 2048, 4096, and 2048 respectively. After the network is trained, the output of the 3rd fully connected layer is taken as the feature of the input hyperspectral pixel.

**Fujitsu Satellite references**


Cai, Z. and Vasconcelos, N., 2018. Cascade r-cnn: Delving into high quality object detection. In Proceedings of the IEEE conference on computer vision and pattern recognition (pp. 6154-6162).

Chen, K., Pang, J., Wang, J., Xiong, Y., Li, X., Sun, S., Feng, W., Liu, Z., Shi, J., Ouyang, W. and Loy, C.C., 2019. Hybrid task cascade for instance segmentation. In Proceedings of the IEEE/CVF Conference on Computer Vision and Pattern Recognition (pp. 4974-4983).

Qiao, S., Chen, L.C. and Yuille, A., 2020. Detectors: Detecting objects with recursive feature pyramid and switchable atrous convolution. arXiv preprint arXiv:2006.02334.

Fu, Y., Zhang, T., Zheng, Y., Zhang, D. and Huang, H., 2019. Hyperspectral image super-resolution with optimized rgb guidance. In Proceedings of the IEEE/CVF Conference on Computer Vision and Pattern Recognition (pp. 11661-11670).

Lin, T.Y., Dollár, P., Girshick, R., He, K., Hariharan, B. and Belongie, S., 2017. Feature pyramid networks for object detection. In Proceedings of the IEEE conference on computer vision and pattern recognition (pp. 2117-2125).

Ren, S., He, K., Girshick, R. and Sun, J., 2016. Faster R-CNN: towards real-time object detection with region proposal networks. IEEE transactions on pattern analysis and machine intelligence, 39(6), pp.1137-1149.

Verma, V., Lamb, A., Beckham, C., Najafi, A., Mitliagkas, I., Lopez-Paz, D. and Bengio, Y., 2019, May. Manifold mixup: Better representations by interpolating hidden states. In International Conference on Machine Learning (pp. 6438-6447). PMLR.

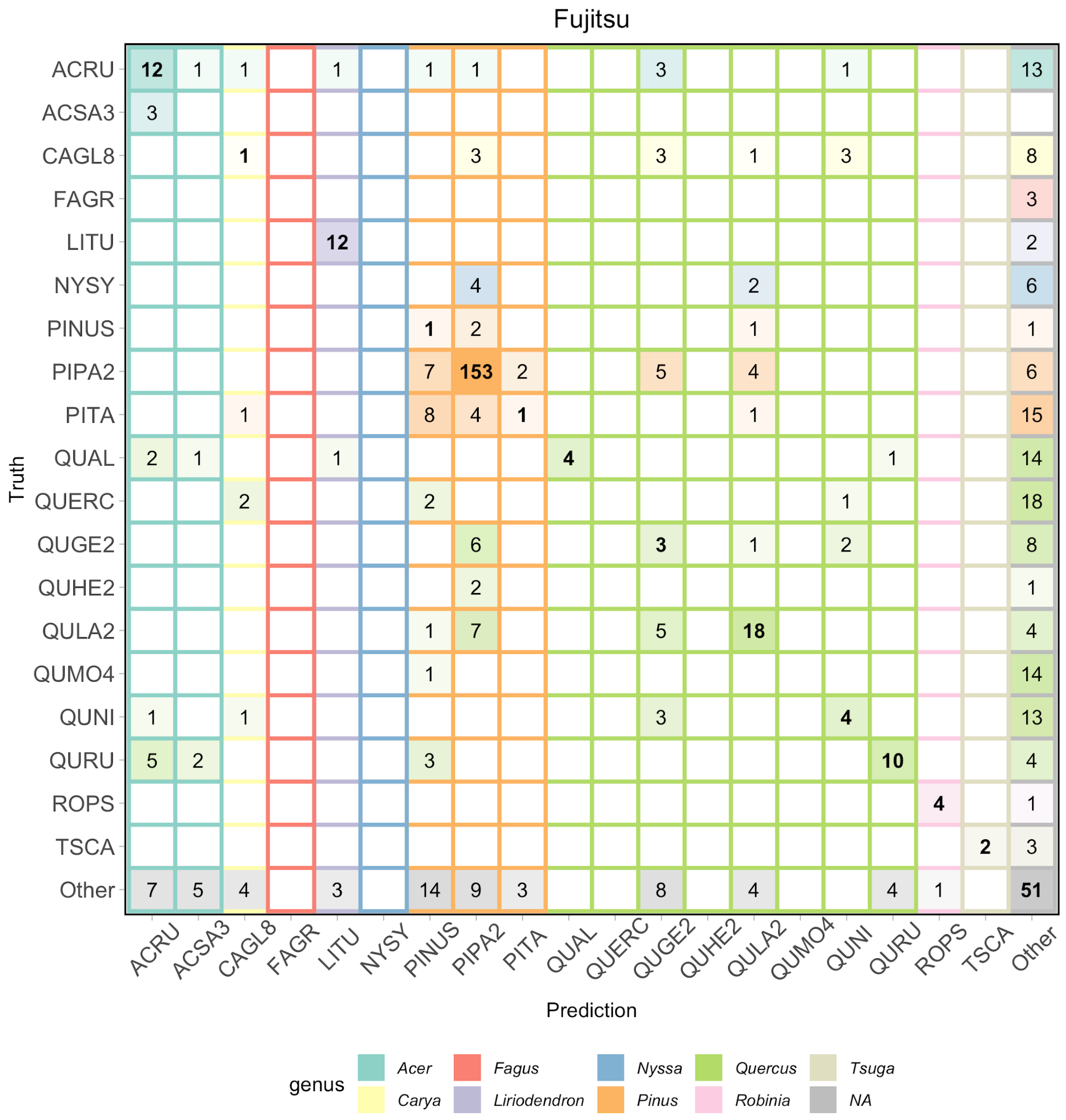


**Figure B3. Confusion matrix for the Fujitsu Satellite team.** Data from all three evaluation sites is included.

### Jeepers Treepers team

**Classification**

We used a two-stage neural network approach to integrate remote sensing data from all three sensors (RGB, hyperspectral, and lidar) and classify individual plant taxa. First we cropped a rectangular RGB image subset for each individual plant using the provided canopy polygons. Using these cropped RGB images, we trained a ResNet convolutional neural network (He et al. 2016) to generate a taxon ID probability vector for each individual crown.

We concatenated this probability vector with the hyperspectral data extracted at crown centroids, along with pseudo-waveform data derived from the lidar point cloud. The pseudo-waveforms contain the density of discrete lidar returns within a series of height bins from a height of zero to the height above ground for any given tree. This concatenated feature vector was passed to a second neural network that learned to fuse the RGB, hyperspectral, and lidar data together. We trained this multimodal neural network (Ngiam et al. 2011), also known as a ”fusion network” (Goodfellow et al. 2016), to minimize a custom “soft F1” loss function, and then post-processed the predictions with a certainty less than 50% from the model by assigning 50% probability to an “other” category.

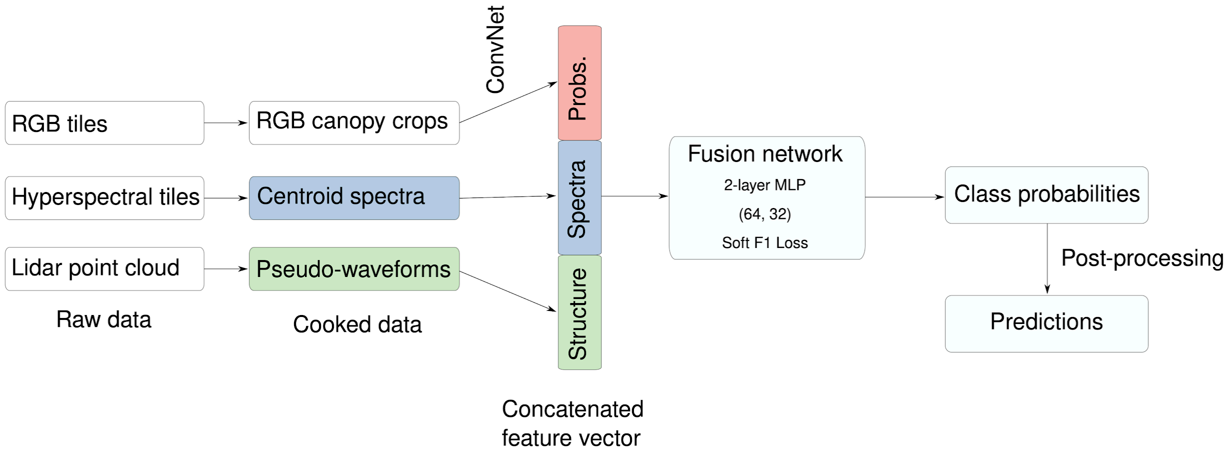


**Figure B4. Classification workflow used by the Jeepers Treepers team.** Raw remote sensing data products were process into formats that describe the spectral and structural characteristics for each individual plant canopy. A pre-trained Convolutional Neural Network (ConvNet) was used to estimate taxon probabilities using the RGB cropped canopy images, and combined these taxon probabilities with hyperspectral reflectance spectra and lidar-derived pseudo-waveforms into a concatenated feature vector. This feature vector was the input to the so-called “fusion network”", a 2-layer multilayer perceptron (MLP) with two hidden layers (size 64 and 32) and trained using a custom “soft F1” loss function, to predict taxon class probabilities for each individual plant. We then applied post-processing including a threshold to assign individuals to an “other” class when the classification confidence was low. Finally, we produced predictions of taxon probabilities.

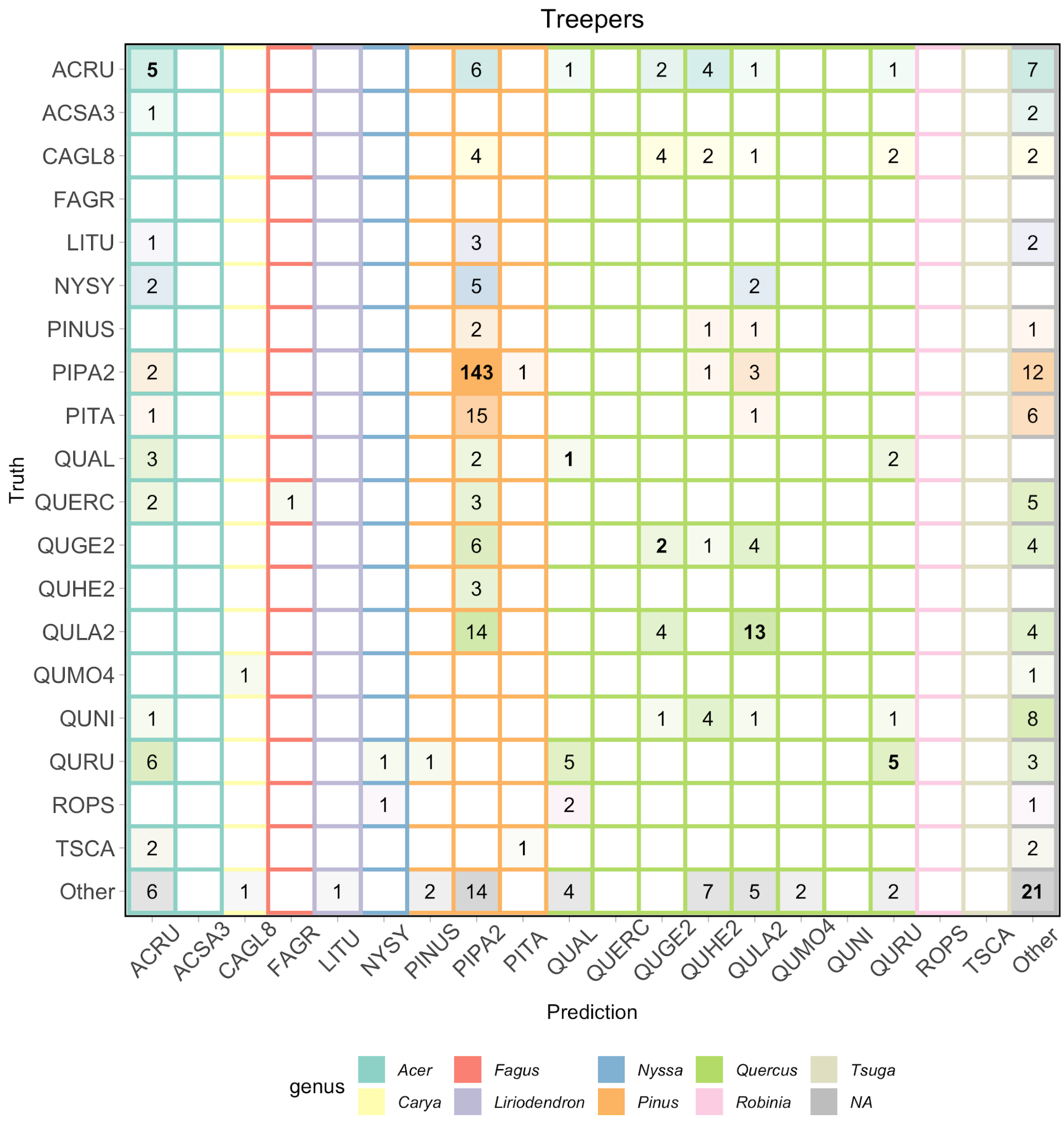


**Figure B5. Confusion matrix for the Jeepers Treepers team.** Data from all three evaluation sites is included.

**Jeepers Treepers references**

He, K., Zhang, X., Ren, S., and Sun, J. (2016). Deep residual learning for image recognition. In Proceedings of the IEEE conference on computer vision and pattern recognition, pages 770–778.

Ngiam, J., Khosla, A., Kim, M., Nam, J., Lee, H., and Ng, A. Y. (2011). Multimodal deep learning. In ICML.

Goodfellow, I., Bengio, Y., Courville, A., and Bengio, Y. (2016). Deep learning, volume 1. MIT press Cambridge.

### Más JALApeñoS team

**Classification**

We used Extreme Gradient Boosting (Chen & Guestrin, 2016) from the 'xgboost' R package for model fitting and prediction (Chen et al., 2015). Our first data processing step was to crop the hyperspectral and canopy height rasters to the ITC bounding boxes of each individual tree in the training set. After removing instances where ITCs spanned 2 remote sensing images, we were left with 1,057 unique tree crowns (648 in MLBS and 409 in OSBS) in our training set. We applied an identical procedure to the testing set (71 ITCs in MLBS, 319 ITCs in OSBS, and 196 ITCs in TALL). Part of the classification challenge was to identify novel species the model was not trained on as 'other' species. To train the model to classify 'other' species, we labeled all individuals from species that had less than 3 individuals in the overall dataset as 'other' species, resulting in 12 individuals falling into this category. We determined the 3 individual thresholds, as with many other parameters used in the modeling process, based on a partial grid search of parameter space (see below for details, Table 1).

Next, we decomposed each cropped hyperspectral and canopy height raster in its corresponding 1 m x 1 m pixels and removed all pixels (n = 367 pixels) with negative values, which likely represent sensor errors. We then removed pixels (n = 2,197 pixels) with a canopy height ≤ 1 m, which may represent bare ground. Additionally, we removed individuals with < 5 pixels from the dataset so the model would not learn from individuals with small amounts of data.

To validate our model, we further split the training dataset into training and validation using a 70/30 split within each species. As such, approximately 70% of individuals within each species and their corresponding pixel-level data were randomly placed into the training set and the remaining 30% were used in the validation set. We conducted the training/validation split at the ITC level to avoid having one ITC being represented in both the training and validation set. In total, we had 23,155 pixels from 721 individuals in the training set and 9,919 pixels from 290 individuals in the validation set.

Following Anderson (2018), we filtered outlier pixels using a PCA-based method. First, we performed a PCA on the 369 hyperspectral bands on the pixels in the secondary training set (n = 23,155 pixels). We then removed a pixel if it was beyond a ± 5 standard deviation (SD) threshold in any of the first 50 principal components. We ran a similar procedure on the validation pixels, using the PCA fit to the secondary training set. In total, this process filtered out 365 pixels from the secondary training set and 235 pixels from the validation set. We then PCA-transformed the hyperspectral imagery data in an attempt to isolate the potentially biologically-relevant components of the hyperspectral imagery that would describe the most variation in reflectance across different species in the dataset.

For Gradient Boosting model parameterization, we set a maximum tree depth of 7 ('max_depth'), a learning rate of 0.1 ('eta'), a minimum child weight of 7 ('min_child_weight'), subsampling the training data by 65% (‘subsample’), and subsampling the prediction data by 50% for each tree (‘colsample_bytree’), and used default values otherwise. Initial model runs were for 5,000 rounds, with the early stopping condition that the model fitting would stop if there was no improvement in model fit for 10 consecutive trees. We ran the final model for a total of 79 rounds, which was determined to be the optimal number of rounds to reduce classification error under 5-fold cross-validation using the ‘xgb.cv’ function.

We then use the fitted Gradient Boosting model to make predictions on each individual pixel in the validation and testing sets, resulting in a matrix containing the probability of each pixel belonging to each species in the training set. To create a species classification prediction at the ITC level, we averaged the species classification probability of each pixel within the ITC for an individual.

For species prediction on the testing set provided by the competition organizers, we followed a similar analysis, except that we did not filter out pixels based on the PCA because that might have removed pixels from species that were not part of the training set.

Finally, to determine the optimal parameter values for the hyperparameters in the Extreme Gradient Boosting model, as well as threshold values for data processing, we used a partial grid search approach. We explored a random subset of the potential range of parameter values (Table B2), and for each parameter combination, fit a Extreme Gradient Boosting model on the secondary training set (n = 23,155 pixels), and calculated the overall species classification accuracy of the fitted model on the validation set. In total, we fit 9,842 models in our partial grid search. The final model we chose to use on the testing set had among the highest values of predictive accuracy on the validation set.

**Table B2. Parameter combinations used in partial grid search to determine optimal parameter values for species classification from hyperspectral data.** ‘Parameter’ is the description of the parameter being varied, ‘Xgboost parameter name’ is the parameter name used in the xgboost function, if applicable, and ‘Range of values’ shows the range of parameter values explored in the partial grid search.

| Parameter | Xgboost parameter name | Range of values |
|---|---|---|
| Threshold for placing into ‘Other’ species | - | 3, 5 |
| Minimum canopy height | - | 1, 3 |
| Minimum pixel count | - | 0, 5 |
| SD threshold for filtering outlier pixels based on PCA | - | 3, 5 |
| Number of PCs to include in model | - | 25, 50, 100, 369 |
| Transform hyperspectral bands with PCA prior to model fitting? | - | True, False |

| | | |
|---|---|---|
| Learning rate | eta | 0.05, 0.1, 0.3, 0.5, 0.9 |
| Maximum tree depth | max_depth | 1, 4, 7, 10, 15 |
| Minimum child weight | min_child_weight | 1, 4, 7 |
| Subsample split of training data in xgboost model | subsample | 0.33, 0.66, 1.0 |
| Subsample split of prediction data in xgboost model | colsample_bytree | 0.2, 0.5, 1.0 |

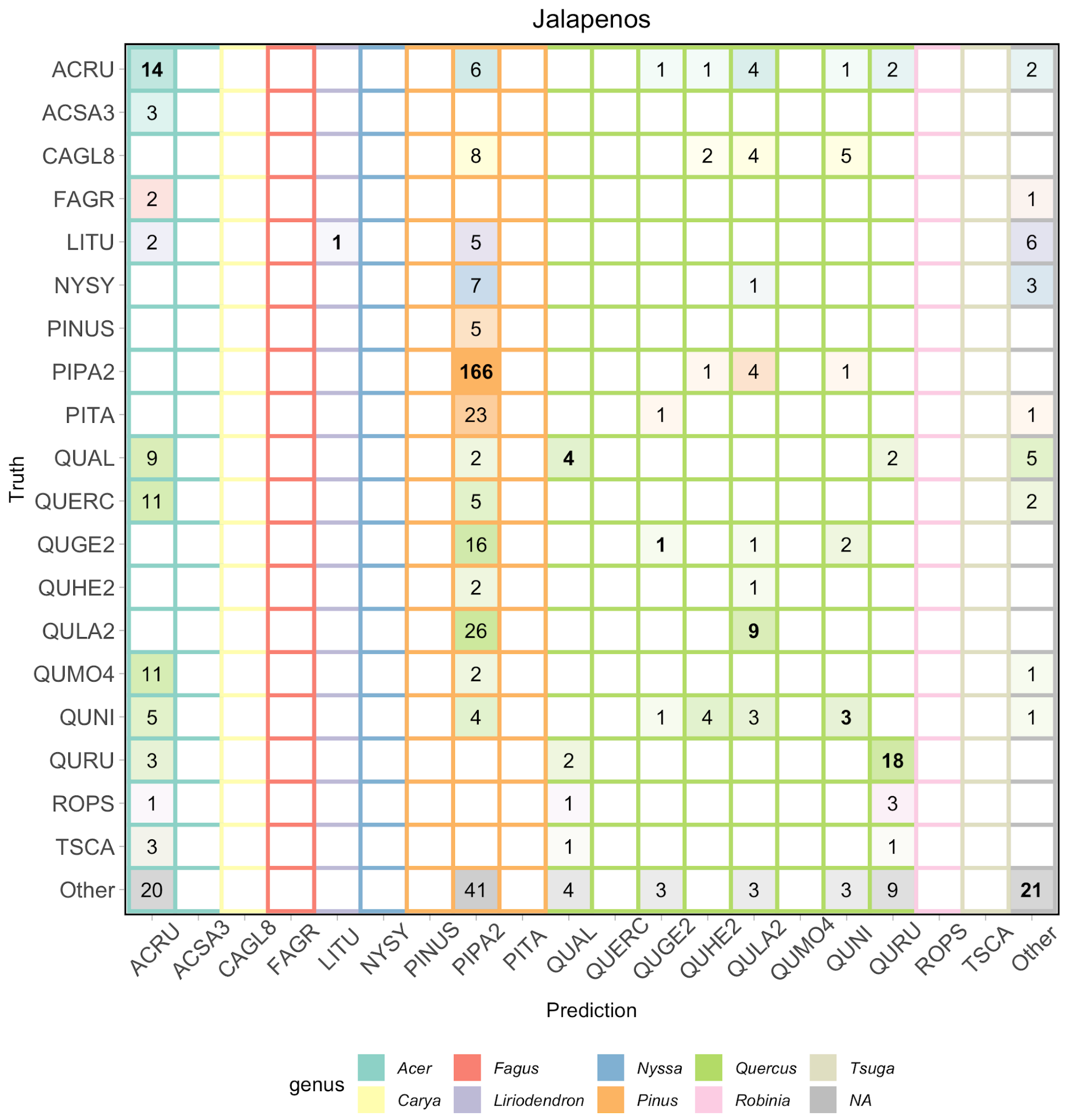


**Figure B6. Confusion matrix for the Más JALApeñoS team.** Data from all three evaluation sites is included.

**Más JALApeñoS References**
Anderson CB. 2018. The CCB-ID approach to tree species mapping with airborne imaging spectroscopy. PeerJ 6:e5666.

Chen T, Guestrin C. 2016. XGBoost: A Scalable Tree Boosting System. In: Proceedings of the 22nd ACM SIGKDD International Conference on Knowledge Discovery and Data Mining. KDD '16. New York, NY, USA: Association for Computing Machinery, 785–794.

Chen T, He T, Benesty M, Khotilovich V, Tang Y. 2015. Xgboost: extreme gradient boosting. R package version 0. 4-2:1–4.

### INRAE-GIPSA team

**Delineation**
We built an adaptive 3D mean shift (AMS3D) algorithm using the LiDAR point cloud data for crown delineation. The approach is based on the strategy published in Tusa et al. 2020. The AMS3D is a nonparametric method using the mode of a probability density function (PDF) to identify clusters of points that may belong to the same object. Specifically, we used the LiDAR point cloud to estimate the PDF based on a kernel profile. The kernel consisted of a weighted moving window used to calculate the mode. Based on the distribution of modes, we defined each individual tree crown (ITC) as the cluster of points converging to the same value. Differently from commonly used mean shift algorithms (Ferraz et al. 2016, Ferraz et al. 2012), our approach was based on adapting a superellipsoid (SE) kernel profile size fitted into an ellipsoid crown model (Fig. B7.). We estimated the parameters for defining the "ellipsoid crown shape model" and the "kernel profile size" based on allometric equations. In particular, we used the slope of the linear regression between the tree height and the crown radius, $m_1$, to derive the radius of the kernel profile circle in the xy-plane, $r_k$; and the slope between the tree height and the crown depth, $m_2$, to adapt the size of the SE kernel profile on the z-axis, $a_k$, We computed the crown radius, $r_t$, in xy-plane and the radius of the intersection of the crown model along the z-axis, $a_t$, to define the ellipsoid crown shape model. Before applying the model, we filtered out all the LiDAR returns lower than 1.5 m, which we associated to ground. The algorithm was initialized for every point and set of parameters $m_1$ and $m_2$.. In cases for which the mean-shift vector did not converge, we used the crown shape model for controlling the kernel size. In that case, the point of maximum height is searched in the cylindrical neighborhood of radius $r_k$ and infinite height to estimate $r_t$ and $a_t$ by equating the SE kernel profile radius $r_k$ to the tree crown radius $r_t$ at half of the tree height. Once the parameters of the ellipsoid crown model are obtained, the radius $r_k$ of the SE kernel profile is derived from the ellipse equation.The cluster points detected as a potential tree were projected into the xy-plane for computing the 2D centroid and for fitting an ellipsoidal crown shape and assigned to a bounding box around the ellipsoidal crown shape.

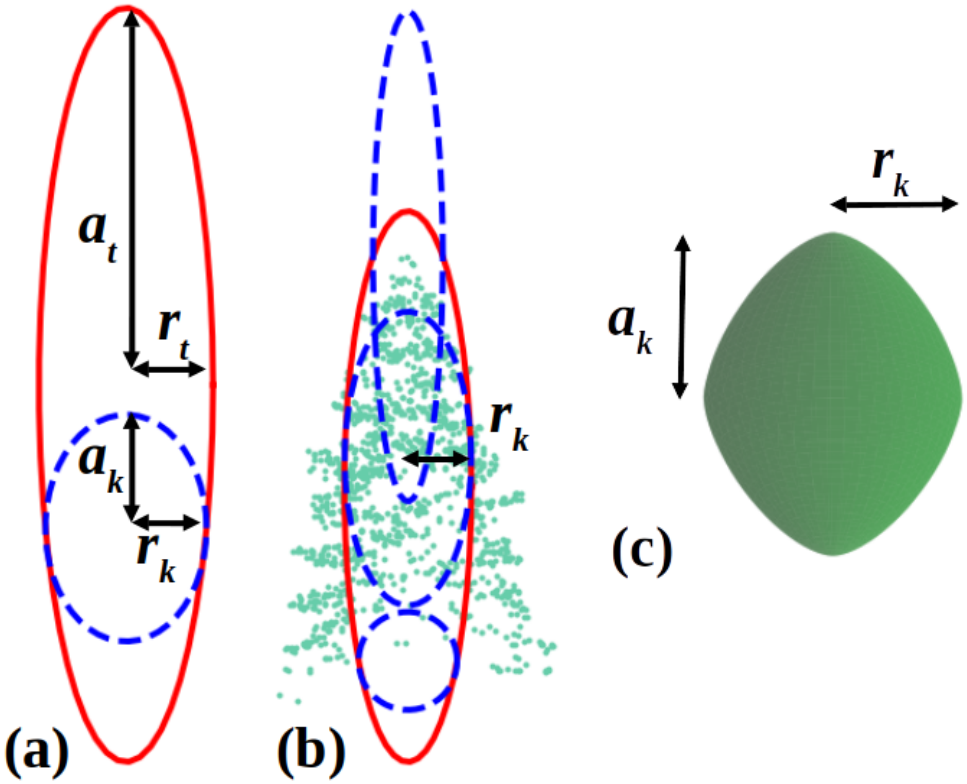


**Figure B7. Kernel profile shape in the AMS3D algorithm.** (a) and (b) Ellipsoid crown shape model in red line and SE kernel profile in dotted blue line. A 3D visualization of the SE kernel profile is shown in (c). In (a), we illustrate the parameters of the ellipsoid crown model ($a_t$, $r_t$) and the SE kernel profile ($a_k$, $r_k$).


Tusa, E., Monnet, J.M., Barré, J.B., Dalla Mura, M., Dalponte, M. and Chanussot, J., 2020. Individual Tree Segmentation Based on Mean Shift and Crown Shape Model for Temperate Forest. IEEE Geoscience and Remote Sensing Letters.

Ferraz, A., Saatchi, S., Mallet, C. and Meyer, V., 2016. Lidar detection of individual tree size in tropical forests. Remote Sensing of Environment, 183, pp.318-333.

Ferraz, A., Bretar, F., Jacquemoud, S., Gonçalves, G., Pereira, L., Tomé, M. and Soares, P., 2012. 3-D mapping of a multi-layered Mediterranean forest using ALS data. Remote Sensing of Environment, 121, pp.210-223.

### Intellisence CAU team

**Delineation**

We use RGB images to train a Faster-RCNN detector. We used a learning rate parameter of 0.02, batch size 8, and ResNet50 as backbone. We used test-Time Augmentation (Shanmugam et al., 2020) in the attempt to improve overall accuracy.

**Classification**

We used a 1D-CNN to pixel values extracted from the hyperspectral data for species classification. The network architecture contains five layers with weights, including the input layer, one convolutional layer, one max pooling layer, the full connection layer, and the output layer. We initially followed two strategies to cope with the imbalanced data: (1) resampling the dataset, and (2) weighting the loss function. Based on the results from validation data, we chose the first strategy for making predictions on the held-out test dataset. To do so, we used a stratified random resampling strategy. For each species, we randomly selected between 400 and 600 individual trees, based on the original sample size of that species. Finally, we filtered outlier pixels using a canopy height model (CHM) threshold to separate pixels belonging to the crown from pixels belonging to the ground. This was particularly important for the OSBS site, where open crowns showed multiple ground pixels within the bounding boxes..

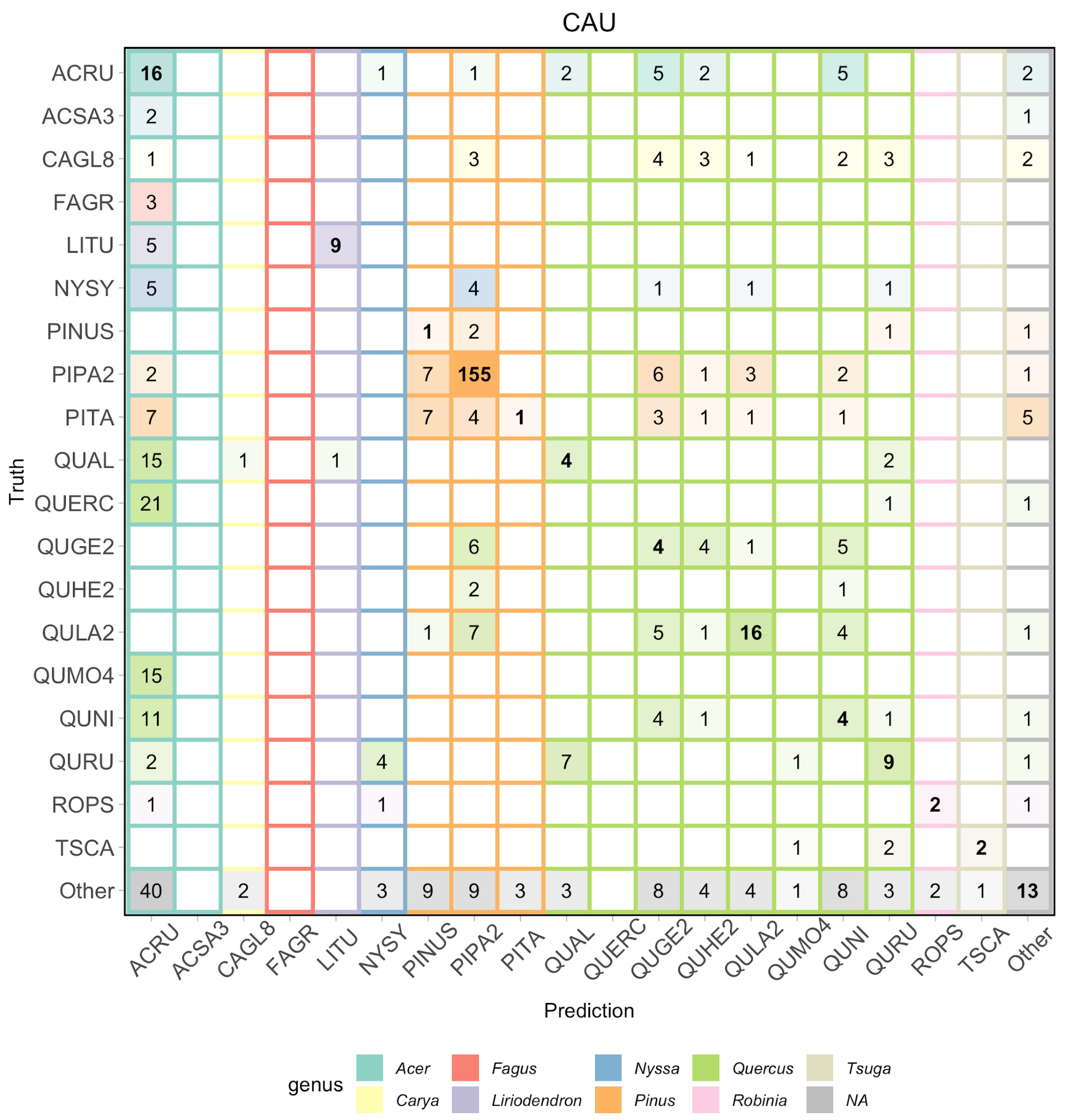


**Figure B7. Confusion matrix for the CAU team.** Data from all three evaluation sites is included.


CAU references

Shanmugam, D., Blalock, D., Balakrishnan, G., & Guttag, J. (2020). When and Why Test-Time Augmentation Works. arXiv preprint arXiv:2011.11156.
